# Deep predictive coding networks partly capture neural signatures of short-term temporal adaptation in human visual cortex

**DOI:** 10.1101/2024.12.06.627148

**Authors:** Amber Marijn Brands, Paulo Ortiz, Iris Isabelle Anna Groen

## Abstract

Predictive coding is a leading theory of cortical function which posits that the brain continually makes predictions of incoming sensory stimuli using a hierarchical network of top-down and bottom-up connections. This theory is supported by prior work showing that PredNet, a deep learning network designed according to predictive coding principles, exhibits several characteristics of neural responses commonly observed in primate visual cortex. However, one ubiquitous neural phenomenon that has not yet been investigated is short-term visual adaptation: the adjustment of neural responses over time when exposed to static visual inputs that are either prolonged or directly repeated. Here, we examine whether PredNet exhibits two neural signatures of temporal adaptation previously observed in intracranial recordings of human participants viewing prolonged and repeated stimuli (Brands et al., 2024). We find that, like human visual cortex, PredNet adapts to static images, evidenced by subadditive temporal response summation: a non-linear accumulation of response magnitudes when prolonging stimulus durations, which results from neurally plausible transient-sustained dynamics in the unit activation time courses. However, PredNet activations also show a systematic response to stimulus offsets, which is absent in the human neural data. For repeated stimuli, PredNet shows slight response suppression for any two images presented in quick succession, but no repetition suppression, a comparatively stronger response reduction for identical than for non-identical image pairs that is robustly observed throughout human visual cortex. We show that these results are stable across multiple training datasets and two different types of loss computation. Lastly, in both PredNet and the neural data, we find a relationship between temporal adaptation and visual input properties, showing that temporally sustained activity is enhanced for more complex scenes containing clutter. All together, these results suggest that the emergent temporal dynamics in the PredNet only partly align with neural data and are linked to low-level properties of the visual input rather than high-level predictions arising from top-down processes.

## Introduction

Predicting the near future is a crucial ability of human perception needed for survival in a dynamic and complex environment. Predictive coding is a prominent theory of brain function and sensory information processing (Rao and Ballard, 1999; Friston, 2005), which postulates that neural circuits learn representations that reflect the statistical regularities of the natural world, signaling deviations from such regularities to higher processing centers. Rao and Ballard (1999) proposed and implemented a hierarchical architecture for predictive coding, referred to as the *predictive coding scheme*, that explains certain important properties of the visual cortex, including endstopping and other extra-classical receptive field effects. This schema has influenced several subsequent works explaining various perceptual and neurophysiological effects (Hohwy et al. 2008; Spratling 2008; Summerfield and Egner 2009; Auksztulewicz and Friston 2016; for a review see Huang and Rao 2011; Friston 2018), while also providing biologically plausible neural dynamics and synaptic update rules (Friston, 2003; Lillicrap et al., 2020; Millidge et al., 2020). These examples suggest that the general idea of predictive coding may be applicable across different brain regions and modalities, providing a promising framework for understanding the structure and function of the neocortex.

Previous studies have attempted to cast the predictive coding scheme proposed by Rao and Ballard (1999) into a modern deep learning framework. One implementation which has been intrinsically designed according to the predictive coding theory is known as PredNet (Lotter et al., 2016, 2020). This deep neural network (DNN) predicts future video frames, whereby each layer in the network makes local predictions in a topdown fashion and computes prediction errors by comparing those predictions to their subsequent input from upper layers. These prediction errors are then in turn fed to subsequent upper network layers, whereby the network learns in a recursive way to construct and update an internal model of its environment. This deep learning architecture approximates human visual learning in two different aspects. First, since the training objective is next frame prediction, there is no need for large amounts of labeled data, as is also the case with visual learning in humans, where just a few or even a single view is often sufficient to achieve robust recognition of the object across a wide range of viewing perspectives, luminance conditions, and contexts (Carey and Bartlett, 1978; Fei-Fei et al., 2006; Lake et al., 2011). Second, PredNet learns representations from video and is therefore optimized to exploit the temporal structure in sequences of images. This is in line with the naturalistic visual environment of biological organisms (including humans) who generally operate in a visual world alive with movement. It is thought that this temporal experience, with objects and scenes undergoing continuous transformations, serves as an important signal for learning about the general structure of the environment, driven by both self-motion of the human viewer and the movement of objects within the scene (Fö ldiák, 1991; Softky, 1995; George and Hawkins, 2005; O’Reilly et al., 2014; Agrawal et al., 2015; Goroshin et al., 2015; Whitney et al., 2016; Schneider et al., 2021). Moreover, due to the naturalistic inputs depicting real-world scenario’s, it is thought that the PredNet is able to exploit temporal regularities in the visual inputs in a manner similar to the human brain. If this is the case, it is also hypothesized that the response dynamics in the network align with those observed in the visual cortex.

Supporting this hypothesis, previous work has demonstrated PredNet’s ability to capture several phenomena related to temporal dynamics observed in visual cortex, including on/off responses, learning effects and perceptual motion illusions (Watanabe et al., 2018; Fonseca, 2019; Lotter et al., 2020; Kirubeswaran and Storrs, 2023). One prominent and ubiquitous property of neural responses that - to our knowledge - has not yet been studied in detail is short-term visual adaptation, which exhibits interesting and complex temporal dynamics, as reported in a series of recent studies on intracranial EEG recordings in human visual cortex (Zhou et al., 2019; Groen et al., 2022; Brands et al., 2024). First, neural responses show subadditive temporal summation, referring to a nonlinear accumulation of response magnitudes when a static visual stimulus is prolonged in time (**Fig. 1A**). Second, neural responses show a reduction in response magnitude when two visual stimuli are shown in quick succession (response suppression), with a stronger response reduction when the two stimuli are identical than when they are different (repetition suppression; **Fig. 1B**). Here, we conduct a new test of the hypothesis of the biological plausibility of PredNet by investigating whether it captures these two neural signatures of temporal adaptation as observed in the human brain. Moreover, previous work has optimized the PredNet by either minimizing prediction errors in only the first, or in all network layers (Lotter et al., 2016; Rane et al., 2020). Minimizing the error in solely the first layer is less realistic from a biological perspective, since it is thought that the brain does not just predict sensory inputs in a single brain area, but instead performs predictions across the visual hierarchy. We therefore also investigate if error minimization across all network layers, which simulates this layered predictive process, leads to more similar temporal dynamics as observed in neural responses.

**Figure 1:**
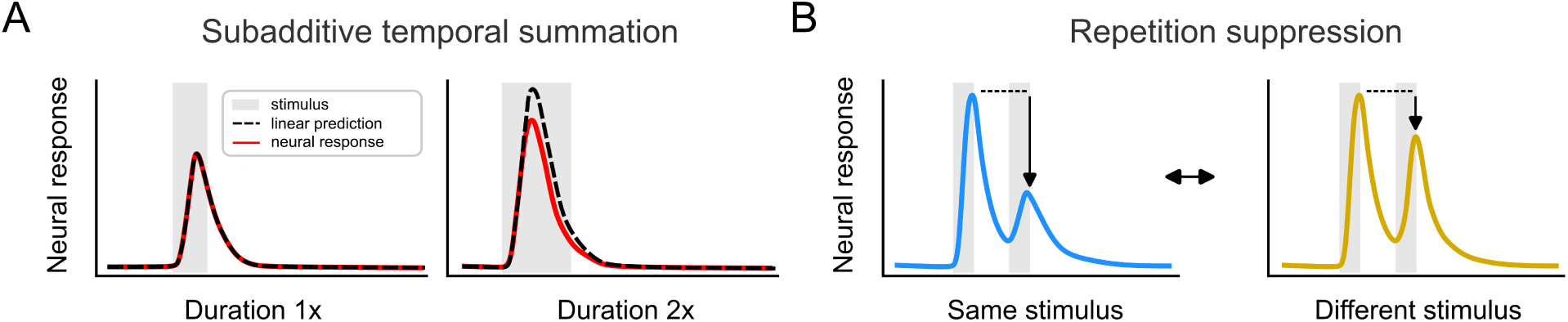
Neural populations show adaptation over time. **A**: When presenting a single, static image, neural responses in human visual cortex exhibit subadditive temporal summation, a nonlinear, compressive accumulation of response magnitudes when prolonging the stimulus duration. Adapted from (Zhou et al., 2019). **B**: When presenting two images separated by a brief gap, responses to the second image are reduced (response suppression); moreover, this response reduction is stronger when the two images are identical, compared to when they are different (repetition suppression).

To test if PredNets trained to predict future video frames in a self-supervised manner exhibit short-term neural adaptation signatures as observed in human visual cortex, we ran an analogous experiment as in our recent human intracranial EEG study (Brands et al., 2024), using the same naturalistic scene images as used in that experiment. To measure temporal summation, we mimicked the presentation of static stimuli with varying durations by presenting PredNet with images for increasing numbers of time steps. To measure response and repetition suppression, we presented PredNet with pairs of images separated by gaps of increasing numbers of time steps, whereby stimulus pairs were either identical (i.e. repeated) or different. We extracted single-unit error activations across all layers and time steps for each stimulus condition, and used these responses to quantify the degree of subadditive temporal summation, response suppression and repetition suppression in the same way as for human broadband iEEG activity in both early and higher visual cortex regions. By comparing the resulting metrics between PredNet and human visual cortex responses, we assess the degree to which PredNet shows brain-like emergent temporal dynamics.

Our analyses yield the following insights. First, PredNet exhibits subadditive temporal summation for single static stimuli, resulting from similar transient sustained-dynamics as observed in the human brain, although it also systematically exhibits offset responses, which are mostly absent in the iEEG data. For repeated stimuli, PredNet exhibits response suppression, but not repetition suppression. Second, computing the loss over all layers during optimization increases the degree of subadditive temporal summation, but does not affect the repetition suppression patterns. Third and last, examination of the unit responses on a trial-by-trial level shows that the sustained activity causing the subadditivity in temporal summation during single stimulus presentations is related to image complexity, with higher responses for cluttered images. Overall, our in-depth comparison of PredNet with human neural recordings shows that PredNet’s emergent temporal dynamics only partly capture temporal adaptation signatures of human visual cortex, suggesting that predictive coding theory may not fully account for neural phenomena related to temporal adaptation.

## Materials and Methods

### Predictive coding networks

#### Background

The original description of the PredNet can be found in Lotter et al. (2016). Briefly, the PredNet consists of a hierarchical stack of layers with each layer *l* containing four different unit types, namely representational (*R_l_*), target (*Â_l_*), prediction (*A_l_*) and error (*E_l_*) units (**Fig. 2A**). At each timestep *t* updating of unit activations occurs through two passes. First, the states of the representational units *R_l_* are updated via a top-down pass, receiving input from both the error units 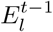 and the representational units 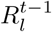 at the same level from the previous timestep, and the representational units 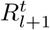 from the layer above at the current timestep, which are first spatially upsampled to match the spatial size of layer *l*. For the representational units a convolutional LSTM is used (Hochreiter and Schmidhuber, 1997; Shi et al., 2015). Following the top-down pass, a bottom-up pass is made where the prediction for the next frame 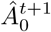 is generated via a convolution of the representational units 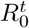. Subsequently, the difference is computed between the actual 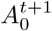 and predicted 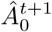 target, consisting of unit-wise subtraction and splitting into positive and negative error populations, yielding the state of the error units for the first layer 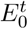. These error units 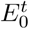 become the input to the next layer *l* + 1 from which a new prediction 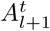 is generated via a convolution, followed by a 2 *×* 2 max-pooling operation.

**Figure 2:**
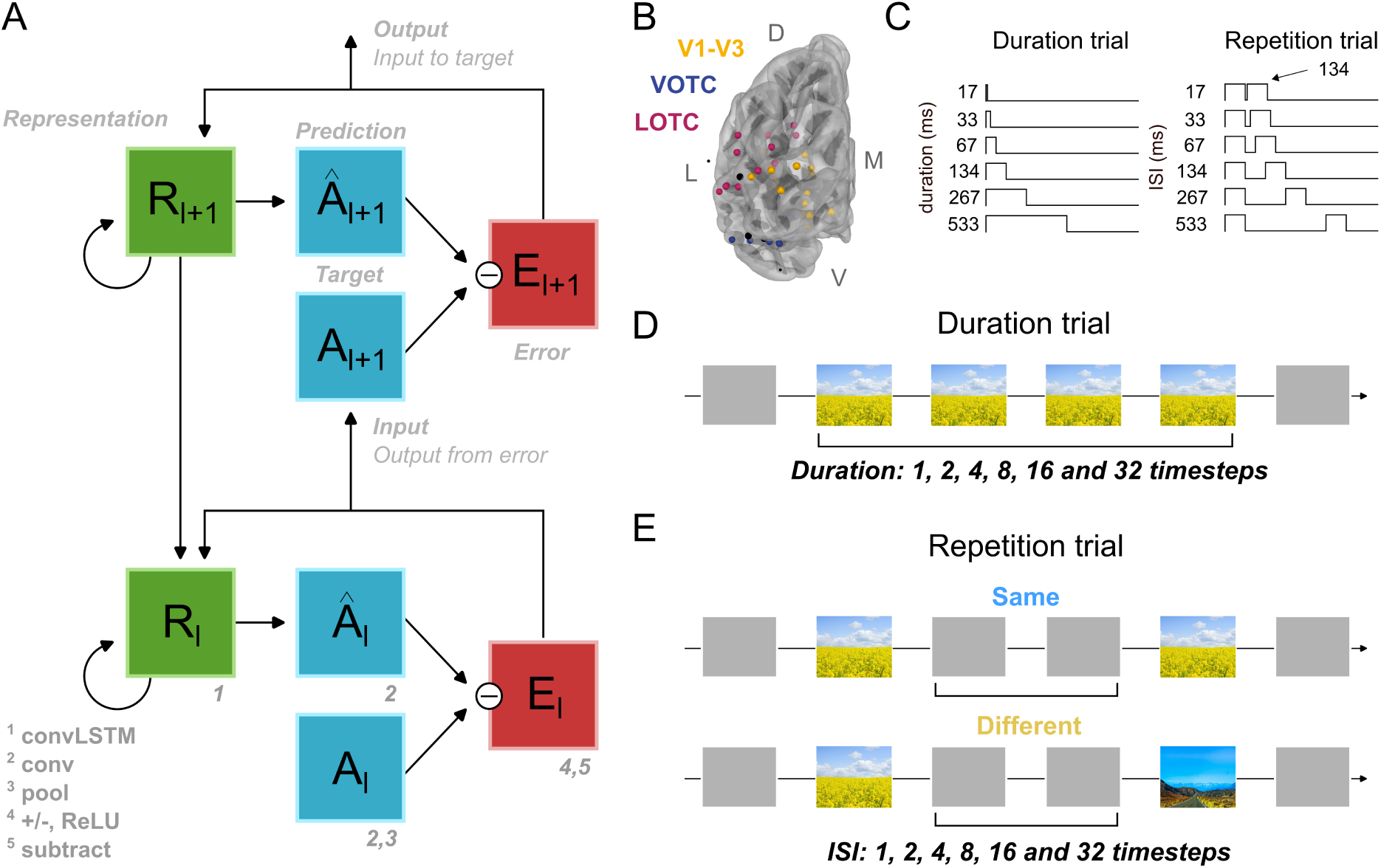
Experimental design. **A**: Information flow within the deep predictive coding network PredNet. Each layer *l* consists of representation units (*R_l_*), which output a layer-specific prediction at each model time step (*Â_l_*), which is compared against a target (*A_l_*) to produce an error term (*E_l_*) which is propagated within the same layer and to the layer above. Figure is adapted from Lotter et al. (2016). **B**: Location of electrodes in the intracranial electrocorticograpy (iEEG) dataset (Brands et al., 2024), depicted on a temporal brain surface (fsaverage; image was created using MNE-Python). Electrodes with robust visual responses were identified in V1-V3 (n = 17), VOTC (n = 11) and LOTC (n = 13). Electrodes not included in the dataset are depicted in black. L = lateral, M = medial, D = dorsal, V = ventral. **C**: Human subjects were presented with visual stimuli whose temporal dynamics were varied in two different trial types. Duration trials (left) consisted of a single stimulus with durations ranging from 17-533 ms. Repetition trials (right) included two stimulus presentations of 134 ms with gaps ranging from 17-533 ms. **D-E**: The PredNet was presented with analogous trial types as the human participants. Single stimulus trials (D) consisted of presentations of a single image for timesteps in powers of two, i.e. 1, 2, 4, 8, 16 and 32. Sequential stimulus trials (E) consisted of the presentation of two images which were either the same (top) or different (bottom) with varying inter-stimulus intervals (same temporal step size as the single stimulus trials).

Formally, given an input sequence *x* (input size of 128 *×* 160 pixels), the units at each layer *l* (with a total of *L* layers) and timestep *t* are updated by an initial top-down pass during which the states of the representational units are computed:

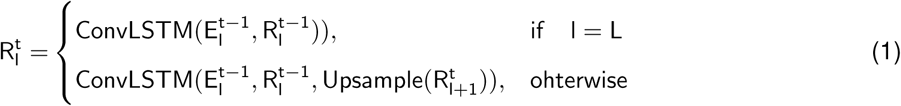

followed by a bottom-up pass during which the states of the predictions, errors and targets are computed:

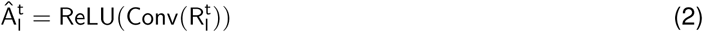

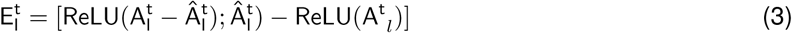

and for *l < L*:

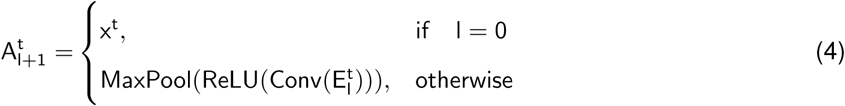

Based on a previously performed hyperparameter search (Lotter et al., 2016), a four-layer model with 3 × 3 filter sizes for all convolutions and stack sizes per layer of 3, 48, 96 and 192 for the *E* and *R* modules was adopted. The model is trained in an unsupervised or ‘self-’supervised manner to perform next-frame prediction, that does not require any external labels or other forms of supervision.

#### Pretrained PredNet

We examined activations derived from a pretrained PredNet which was written in Keras and optimized on 10-frame sequences of 128 *×* 160 pixel RGB videos from the KITTI dataset (Geiger et al., 2013), which consists of a collection of videos obtained from a car-mounted camera while driving in Germany. Sequences were sampled from the “City”, “Residential” and “Road” categories, with 57 recording sessions used for training. Code to run and train the model on the KITTI dataset is available at https://github.com/coxlab/prednet.git and methodological details can be found in Lotter et al. (2016) and Lotter et al. (2020).

#### Datasets and training procedures

To test the robustness of the results of the pretrained network, we also trained a number of PredNet instance from scratch on the following four datasets: three video’s belonging to the “Walking Tours” dataset (Venkataramanan et al., 2023). The “Walking Tours” dataset contains a set of first-person hours-long videos, captured in a single uninterrupted take, depicting a large number of objects and actions with natural scene transitions. We selected two videos recorded in urban areas, including *Amsterdam* and *Venice*, and one video from a *wildlife* safari. All four videos were preprocessed, including downsampling to 15 frames-per-second and resizing to 128 *×* 160. For detailed info about each video, see **Supplementary Table 1**.

Each network was trained for 150 epochs on one of the videos, with each epoch consisting of 500 samples. Samples consisted of a series (batch size of 4) of 10-frame sequences which were randomly selected. Mean squared pixel error loss was computed (see below) over all time steps using the Adam optimizer with learning rate of 0.001, reduced by a factor of 0.9 each 100 batches. Model weights were updated after each batch and training frames within one sequence were processed in order. For a controlled comparison, we also retrained one PredNet on the KITTI dataset mentioned above. To assess reliability, multiple instances (n = 3) with different random initial weights were trained for each video dataset.

#### Computation of the loss

To investigate how the error minimization during training influences the emergent temporal dynamics across network layers, we optimized the PredNets using two different objective functions. More specifically, the training loss is formalized as follows for a model of *L* layers and *T* timesteps:

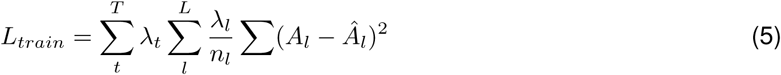

where *λ_t_* are the weighting factors by time (*λ*_0_ = 0, *λ_t>T_* = 1), *λ_l_* are the weighting factors by layer and *n_t_* the number of units in the *l*th layer. We applied two different weighting schemes, with the loss *L_train_*concentrating either on i) solely the lowest layer (i.e. *λ*_0_ = 1, *λ_l>_*_0_ = 0), from here on referred to as *L*_0_ or ii) the loss was moderately enforced by upper layers as well (i.e. *λ*_0_ = 1, *λ_l>_*_0_ = 0.1), from here on referred to as *L_all_*. Unless stated otherwise, the *L*_0_ loss was used for optimization for all analyses.

### Human brain recordings

To determine to what degree emergent temporal dynamics in the PredNet exhibit the same neural adaptation signatures as observed in neural responses in visual cortex, we reanalyzed a dataset from Brands et al. (2024). In this study, iEEG data were collected from four participants implanted with subdural electrodes for clinical purposes at the New York University of Grossman School of Medicine, who were presented with two trial types, which are explained in more detail below. Participants were implanted with standard clinical strip, grid and depth electrodes. Raw voltage time courses were referenced to the common average for each electrode strip, and then filtered into separate 10Hz wide frequency bands ranging between 50-200 Hz. This was followed by calculating the power envelope of each band-pass filtered time course which were then averaged across bands to yield an time-varying broadband time course. By aggregating responses across patients, we obtained 41 visually response electrodes which we separated into one lower-level group (V1-V3, *n* = 17) and two higher-level groups, covering ventral-occipital cortex (VOTC, *n* = 11) and lateral-occipital cortex (LOTC, *n* = 13) using retinotopic atlases (**Fig. 2B**). For more details regarding data processing and electrode localization and selection, see Brands et al. (2024).

### Stimuli

For the experimental setup, PredNet instances were presented with the same stimuli that were used during collection of the neural dataset described above (Brands et al., 2024). This dataset consisted of 288 images (569 *×* 568 pixels) belonging to one of six categories, including buildings, bodies, faces, objects and scenes. All categories except scenes however consisted of the visual category depicted on a gray background, while PredNets are optimized on image sequences of video frames covering all pixels. To minimize potential effects of input distribution shifts on PredNet performance and activations, we therefore only used the 48 images from the scene category that did not contain a gray background. These images consisted of indoor, outdoor man-made and outdoor natural scenes (16 images each). This stimulus set was used for all comparisons of temporal dynamics between PredNets and neural recordings. For our trial-by-trial analyses (see below), we additionally used a larger image dataset collected by Groen et al. (2013) consisting of 1600 scene images (640 *×* 480 pixels) of a wide variety of indoor and outdoor places, landscapes, forests, cities, villages, roads, images with and without animals, objects, and people. The images were selected such that one half of the set contained mostly man-made, and the other mostly natural elements.

### Experimental design

To compare the emergent temporal dynamics in the PredNet with those in the iEEG dataset, we emulated the stimulus conditions from the human experiment and presented the network with two different trial types. In the human experiment, these two trial types were referred to as duration and repetition trials (**Fig. 2C**). Duration trials showed a single stimulus for one of six durations (**Fig. 2C**, *left*), namely 17, 33, 67, 134, 267 and 533 ms. Repetition trials contained a repeated presentation of either the same or two different images with fixed duration (134) but variable inter-stimulus interval (**Fig. 2C**, *right*), ranging between 17-533 ms with the same temporal step sizes as the duration trials. The PredNets were also presented with duration (**Fig. 2D**) and repetition (**Fig. 2E**) trials, whereby the stimulus duration and inter-stimulus interval were varied across six different temporal conditions defined in powers of two, i.e. 1, 2, 4, 8, 16 and 32 model time steps. An example sequence for each trial type and temporal condition is illustrated in **Supplementary Figure 2**. Each trial consisted of 45 model timesteps and the pixel values of inputs at model timesteps which did not contain an image were set to 0.5.

### Summary metrics

To characterize the temporal dynamics of the PredNets’ error unit activations and the iEEG broadband responses we computed several summary metrics, which will be explained in more detail below. We focus on the error rather than the representation (*R*) or target (*A*) units because error unit activations were previously found to exhibit the highest similarity with neural data (Lotter et al., 2020).

#### The degree of subadditive temporal summation

To determine the degree of subadditive temporal summation of either the neural or PredNet responses with increasing stimulus durations, we fitted the response magnitude across durations with a logarithmic:

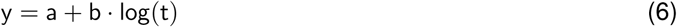

or linear:

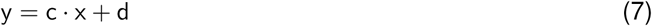

objective function, where *y* is the response magnitude, *x* the stimulus duration and [*a, b*] and [*c, d*] are free parameters for the logarithmic and linear function, respectively. To then quantify the degree of subadditivity, we computed the ratio between the coefficient of determination (*R*^2^) for the logarithmic and linear fit, whereby *R*^2^ *<* 1 suggests that temporal summation is linear, while *R*^2^ *>* 1 indicates that the summation of responses is subadditive.

#### The recovery from adaptation for repeated stimuli

To quantify the degree of adaptation to repeated stimuli, we calculated the response magnitude of the second stimulus as a proportion of the first. Measuring the degree of recovery for the neural responses, however, is not straightforward because the response to the first stimulus overlaps with the second due to lingering activity of the first response after stimulus offset, particularly for shorter ISIs. To address this, we averaged the response time courses for the 134 ms duration stimulus for single trials and all the repeated stimulus trials up to the onset of the second stimulus, thereby obtaining an average of the response to the first stimulus. This average was then subtracted from the repeated stimulus trials to isolate the second response. Recovery from adaptation was then defined as the Area Under the Curve (AUC) of the second response relative to the AUC of the first. For the PredNet activations, we used a similar approach, whereby we subtracted the first response, obtained from a duration trial with one model timestep, from the repeated sequence trial, after which we computed the ratio between the first and second response (AUC of the second response/AUC of the first response).

To then describe the degree of recovery from response suppression both for the neural responses and PredNet responses, we fitted the recovery values across temporal conditions with the logarithmic function(**Eq. 6**). Moreover, to determine whether responses exhibit repetition suppression, we averaged the recovery values across ISIs separately for the trials showing either the same or different images pairs.

#### Sustained level activity

To investigate the relationship between the degree of subadditivity during duration trials and image complexity, we determine the magnitude of the sustained response magnitude for the longest stimulus duration relative to the transient response magnitude (maximum/peak). For the PredNet activations, we obtain the level of the sustained magnitude by determining the activity for the timestep prior to stimulus offset. For the neural responses, the sustained magnitude of neural responses is more challenging to determine due to noise, which obscures the transitions at stimulus offset. To obtain an estimate of the sustained level, we therefore averaged the neural response magnitudes from [-100, 0] relative to stimulus offset.

#### Low-level image statistics

To relate the PredNets’ temporal dynamics to the complexity of the visual input, we performed a trial-by-trial analysis, where for each image we computed a set of low-level contrast statistics, motivated by a series of previous studies that have linked neural activity to scene complexity (Scholte et al., 2009; Groen et al., 2013, 2018, 2012a,b). Specifically, we used a biologically-plausible image-computable model of scene clutter that simulates the response distribution of isotropic, contrast-selective receptive fields in the magno- and parvo-cellular pathways in the LGN. Based on the estimated responses distributions, the model extracts two summary parameters per image, namely contrast energy (CE) and spatial coherence (SC) which have been shown to modulate response amplitudes in EEG (Groen et al., 2013). CE is calculated as the distribution mean, thus indicating with average local contrast strengths in the image, while SC is calculated as the mean divided by the standard-deviation of the distribution, thus indicating the relative spread in local contrast across the image. Code for calculating CE and SC for a give input image is available at https://github.com/irisgroen/LGNstatistics.

### Data and code availability

All code used for the purpose of this paper can be found at the GitHub repository https://github.com/ABra1993/nAdaptation_PredNet.git and the iEEG data can be found at https://openneuro.org/datasets/ds004194/versions/2.0.0.

## Results

Here, we investigate to what degree a PredNet optimized for next-frame prediction (**Fig. 2A**) exhibits sub-additive temporal summation and repetition suppression, two signatures of short-term temporal adaptation observed in human visual cortex. First, we show results for a pretrained PredNet presented with single (**Fig. 2D**) and repeated (**Fig. 2E**) naturalistic images with variable stimulus duration and inter-stimulus interval (ISI), and examine to what degree the emergent network dynamics align with temporal dynamics of iEEG broadband time courses recorded in both early and higher level visual areas (Brands et al., 2024). Next, we show how layer-specific error minimization during optimization influences the emergent temporal dynamics across network layers and the correspondence with neural data. Finally, we perform a trial-by-trial analysis and link differences in temporal dynamics on the individual image level to scene complexity.

### Neural responses in human visual cortex exhibit subadditive temporal summation

We first describe temporal dynamics in responses to duration-varying stimuli in the iEEG broadband data. **Figure** 1 illustrates that neural timecourses exhibit subadditive temporal summation, which refers to the phenomenon that the additional visual exposure resulting from longer presentation durations does not accumulate into a linearly increasing neural response. As described in Groen et al. (2022) and Brands et al. (2024), this subadditivity results from transient-sustained dynamics in the neural response time course, which can be clearly seen in **Figure 3A** (*top*). Neural responses to visual stimulus onsets show an initial transient, which for short stimulus durations is the only part of the response. As stimulus duration increases, this transient response saturates, and a lower-amplitude sustained response emerges. Because prolonging stimulus duration no longer increases the peak transient, and only adds more of the relatively lower-amplitude sustained component, the overall response to the image grows progressively less with longer stimulus durations (**Fig. 3B**). This compressive effect is evidenced by a qualitatively better fit for a logarithmic than a linear function between stimulus duration and summed response magnitude in V1-V3 (dependent-samples T-test computed over the electrodes, *t*_(16)_ = −5.41, *p <* 0.001) and LOTC (*t*_(12)_ = −6.23, *p <* 0.001), with a similar pattern in VOTC (**Fig. 3C**). Moreover, we find a significant relation between the difference in explained variance for these log versus linear fits over the duration-wise summed response and the ratio of the transient and sustained response magnitude (linear regression, *R*^2^ = 0.13, *p* = 0.021; **Supp. Fig. 2A**), demonstrating that the subadditivity of neural response magnitudes is associated with the proportion of transient to sustained activity. To conclude, these results highlight a robust signature of short term-temporal adaptation in human visual cortex: when stimuli are viewed for longer durations, neural activity accumulates non-linearly, as a result of transient-sustained response dynamics in the response time course.

**Figure 3:**
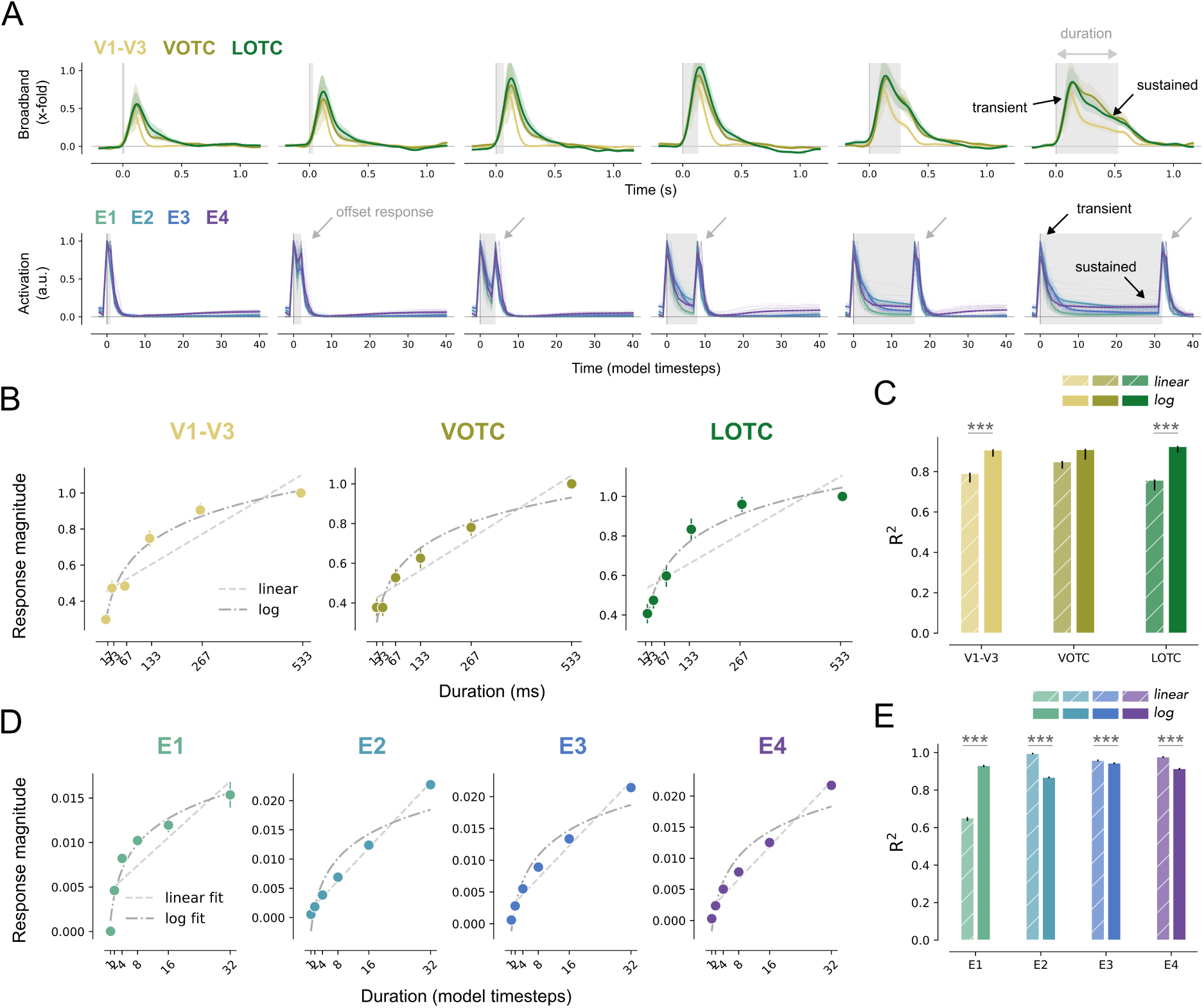
Neural responses and PredNet both exhibit subadditive temporal summation for single image presentations. **A**: *Top*, Normalized broadband iEEG responses for electrodes assigned to V1-V3 (n = 17), VOTC (n = 11) and LOTC (n = 13) to single stimuli (gray). Responses are shown separately per stimulus duration from shortest (17 ms) to longest (533 ms, right). Response shapes are characterized by a transient response (peak) and, for longer stimulus durations, a sustained response (here measured as response magnitude at stimulus offset). *Bottom*, PredNet activations of the error units for the same temporal conditions separately per network layer (i.e. E1, E2 E3 and E4). Similar to neural iEEG responses, Prednet activations show a sustained response for longer but not shorter stimulus durations. Unlike neural responses, PredNet units additionally exhibit a prominent offset response (gray arrows). **B**: Sum of broadband response time courses between 0 and 1 s after stimulus onset for each stimulus duration (circle markers) for V1-V3 (left), VOTC (middle) and LOTC (right) normalized to the longest duration stimulus (533 ms). The lines are fitted to the response magnitudes using either a linear or logarithmic function (see Materials and methods, *Summary metrics*). **C**: Explained variance (coefficient of determination) of summed response magnitude per condition by a linear (striped) or logarithmic (solid) curve for V1-V3 (yellow), VOTC (pink) and LOTC (blue). **D-E** Same as B-C but for the error units of the PredNet model, separately for each model layer. ^∗^ p *<* 0.05, ^∗∗^ p *<* 0.01, ^∗∗∗^ p *<* 0.001.

### Error units in the first layer of PredNet capture subadditive temporal summation

To determine whether the PredNet also exhibits subadditive temporal summation, we emulate the iEEG experiment by presenting the network with the same duration-varying stimuli (see Materials and Methods, *Experimental design*). Similar to the iEEG broadband responses, the PredNet also shows transient-sustained dynamics (**Fig. 3A**, *bottom*): error units across all layers show a transient response at stimulus onset, with a sustained response emerging as the stimulus duration increases. We quantify the degree of temporal summation of the unit responses the same way as for the neural data (**Fig. 3D**) and find the strongest subadditivity for units in the first network layer (*E1*), while higher-level layers show comparatively less sub-additive summation. Indeed, in *E1*, the increase in response magnitude with stimulus durations is best fit with a logarithmic function (dependent-samples T-test computed over the images, *t*_(47)_ = −21.92, *p <* 0.001), while for layers *E2-4* a linear function fits better (E2, *t*_(47)_ = 56.45, *p <* 0.001; E3, *t*_(47)_ = 3.51, *p <* 0.001; E4, *t*_(47)_ = 13.72, *p <* 0.001; **Fig. 3E**). In line with the neural data, we also find a significant relation between the degree of subadditive summation and ratio of the transient and sustained response magnitude in both the first and higher PredNet layers (linear regression, *R*^2^ = 0.89, *p <* 0.001; **Supp. Fig. 2B**).

Notably, we also observe a strong discrepancy between PredNet unit activation timecourses and broadband iEEG responses: PredNet shows a second activation peak to the offset of the stimulus which is absent in the average neural response time courses. As discussed in Brands et al. (2024), offset responses were indeed rarely observed in this iEEG broadband dataset, as well as a different iEEG dataset which measured temporal dynamics of visual cortex responses in a larger number of participants (Groen et al., 2022). Another iEEG study analyzing a single participant did find offset responses but mostly in a subset of electrodes that were specifically tuned to the peripheral visual field (Zhou et al., 2019). We will return to the discrepancy regarding offset responses in PredNet versus human visual cortex in the Discussion. Overall, these results show that the PredNet accurately captures subadditive temporal summation in neural responses, with error units in the first layer exhibiting the strongest subadditivity across stimulus durations.

### Unlike visual cortex, PredNet does not exhibit short-term repetition suppression

So far, we found that PredNet captures subadditive temporal summation for duration-varying stimuli. Another neural signature of temporal adaptation commonly observed in human visual cortex is a response reduction when stimuli are repeated. In our neural iEEG dataset, broadband responses to a second stimulus shown shortly a first one are indeed reduced (**Fig. 4A**, *top*). This suppression is strongest for shorter inter-stimulus intervals (ISIs) and gradually recovers as the gap between two stimuli increases. Moreover, this response reduction is stronger when the second stimulus is the same as the first, compared to when it is different, a phenomenon known as repetition suppression (RS). We quantified the level of response suppression for each ISI and repetition condition by computing the Area Under the Curve (AUC) of the broadband time courses for the second stimulus, divided by the AUC of the response to the first stimulus (**Fig. 4B**; see Materials and Methods, *Summary metrics*). A value less than 1 here indicates a response suppression (and hence adaptation), while a value of 1 or higher would indicate recovery from adaptation or even response enhancement. We observed response suppression across all ISIs, which was substantially stronger for same compared to different stimuli in all visual areas (dependent-samples T-test computed over the electrodes, V1-V3, *t*_(16)_ = −8.17, *p <* 0.001; VOTC, *t*_(10)_ = −5.09, *p <* 0.001; LOTC, *t*_(12)_ = −3.21, *p* = 0.008; **Fig. 4C**). This is consistent with a large body of literature reporting RS in both human and non-human subjects, across many visual areas, and experimental paradigms (for a review, see Grill-Spector et al. 2006).

**Figure 4:**
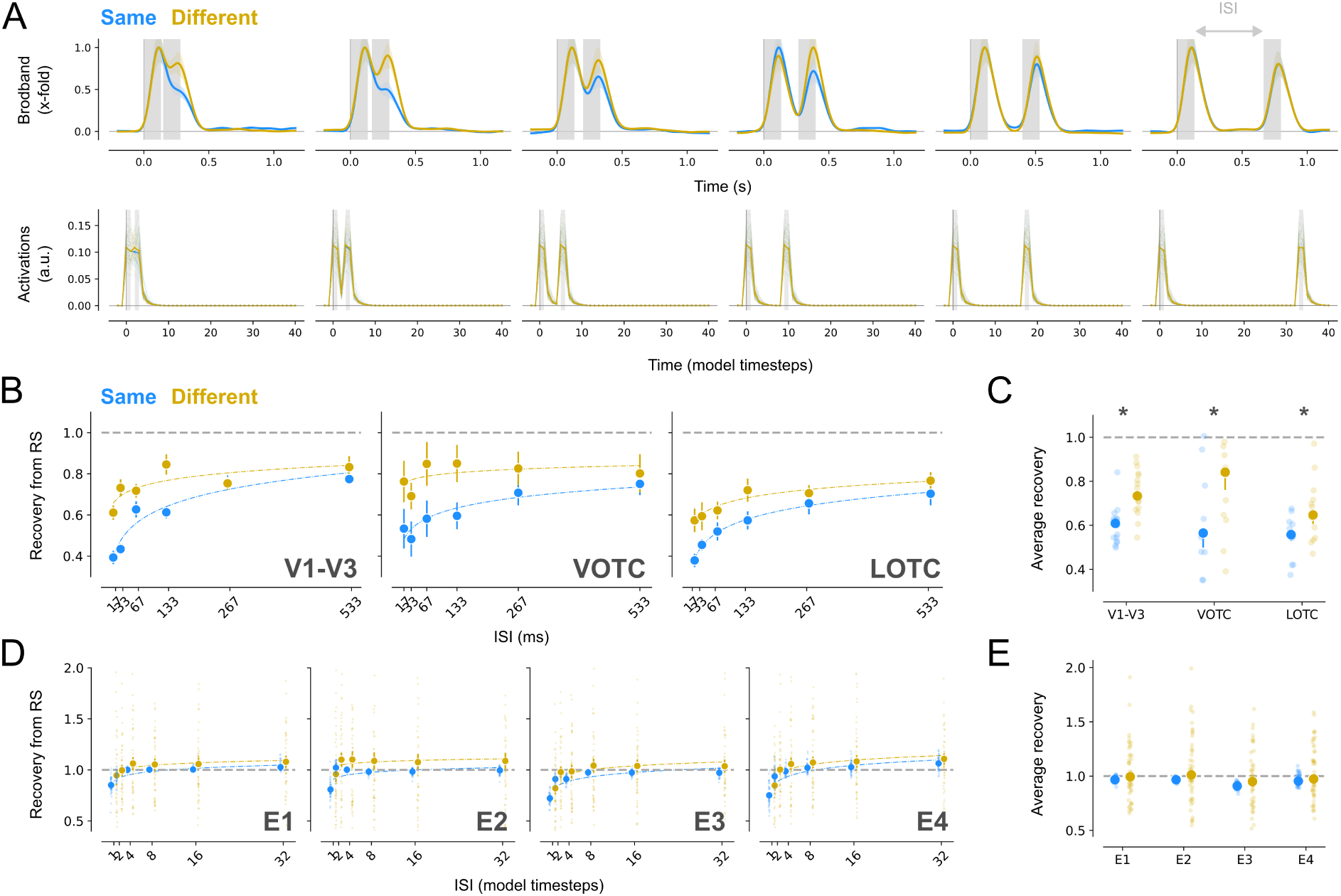
PredNet does not exhibit stronger suppression for same compared to different inputs shown in sequence. **A**: *Top*, Normalized broadband responses for electrodes assigned to V1-V3 (n = 17), VOTC (n = 11) and LOTC (n = 13) to repeated visual stimuli (gray). Responses are shown separately per ISI from shortest (17 ms, left) to longest (533 ms, right). The reduction in response of the second stimulus recovers with increasing ISI. *Bottom*, PredNet activations of the error units in the first network layer for analogous temporal conditions. **B**: Recovery from adaptation for V1-V3 (left), VOTC (middle) and LOTC (right), computed as the ratio of the Area Under the Curve (AUC) between the responses to the first and second stimulus. The fitted curves express recovery as a function of the ISI. The dotted grey line depicts a recovery of 1 (i.e. when the magnitude of the first and second response is the same). **C**: Average recovery from adaptation for each area In all areas, response suppression is stronger for same compared to different stimuli (repetition suppression). **D-E**: Same as B-C for the error units of the PredNet plotted separately per network layer (i.e. E1, E2 E3 and E4 for layer 1 to 4, respectively). The PredNet fails to capture repetition suppression. ^∗^ p *<* 0.05.

In comparison, PredNet units show some response suppression for short ISIs, but unit activity quickly recovers as the number of model steps between the first and second stimulus increases (**Fig. 4A**, *bottom*). This recovery occurs comparatively much faster than for neural responses, which still exhibit substantial suppression for the longest ISI; in contrast, we already observe near-full recovery across all PredNet layers for the second-to-shortest ISI (**Fig. 4D**). Moreover, PredNet does not exhibit the hallmark signature of repetition suppression, i.e. the prominent difference between same and different repeated stimuli as seen in neural data. For the shortest ISIs, the unit activations are slightly lower for same than different image pairs, but the response reduction for longer ISIs is remarkably similar, across all network layers (**Fig. 4D**). Interestingly, the absence of response suppression in PredNet is very stable across same image pairs, while different image pairs exhibit a large spread around zero, showing both positive and negative recovery values (**Fig. 4E**). This suggests that in some cases, the a different preceding image induces a suppression of error unit activity, but in other cases it leads to an enhancement; while for same pair images, there is neither suppression nor enhancement. This lack of suppression is surprising given the *a priori* expectation that same image pairs should be more predictable to PredNet (resulting in lower error unit activations). Importantly, this pattern is again different than was found in the neural data, where responses showed similar variability in response suppression for both same and different image pairs (**Fig. 4C**).

One potential explanation for the modest suppression and the lack of RS in PredNet is that we presented stimuli for just one model timestep, whereby units did not have the opportunity to update their predictions during a longer exposure to the first image. As shown in **Fig. 3B** and discussed above, PredNet error units start to exhibit pronounced offset responses when exposed to longer stimulus durations (i.e., multiple time steps); the observed transient-sustained dynamics also indicate that consecutive repeated exposure to the same image - without presenting grayscale image in the ISI - results in reduced error unit activity, which is then followed by an abrupt response increase at offset due to the presentation of a (presumably unpredicted) grayscale screen. To determine if PredNet exhibits stronger response suppression and repetition suppression for more prolonged exposure conditions, we also ran the repetition experiment using durations of 8 time steps per stimulus (**Supp. Fig. 3**). However, this analysis yielded similar results: PredNet units already exhibited near-full recovery from adaptation for relatively short ISIs, although interestingly, layers *E3-4* showed slightly more suppression for longer ISIs. Notably, the overall degree of recovery was again similar for same and different image pairs, indicating a lack of RS. All together, these findings demonstrate that PredNet shows less temporal adaptation for repeating stimuli compared to our neural recordings, and notably does not capture the strong repetition suppression effects observed in human visual cortex.

### Consistent emergent temporal dynamics when optimizing PredNet on various datasets

Results so far show that units in the first layer of a pretrained PredNet exhibit subadditive temporal summation, while none of the layers’ units exhibit repetition suppression. To assess the robustness and generalizability of these findings, we trained several new PredNet instances on four different datasets, specifically the *KITTI* dataset (Geiger et al., 2013) and three videos belonging to the “Walking Tours” dataset (Venkataramanan et al. 2023; see Materials and Methods, *Training procedures*); the first two videos contained footage of urban areas (*Amsterdam* and *Venice*) and the third was recorded during a *wildlife* safari. We chose datasets derived from walking tour footage because they contain more diverse and also more biologically plausible inputs than the *KITTI* dataset, since videos were collected from the perspective of a walking human, as opposed to a car-mounted camera. Training was successful for each dataset, indicated by the fact that all losses converged well before the end of training (**Supp. Fig. 4A**), and sample predictions demonstrate that our in-house trained networks were able to make accurate predictions on the dataset they were trained on, such as trajectory of passing cars and shadows on the road (*KITTI*, **Fig. 5A**), sliding motion of the camera (*Venice*, **Fig. 5B**), approaching persons on a pedestrian road (*Amsterdam*, **Fig. 5C**) and animal movements (*wildlife*, **Fig. 5D**).

**Figure 5:**
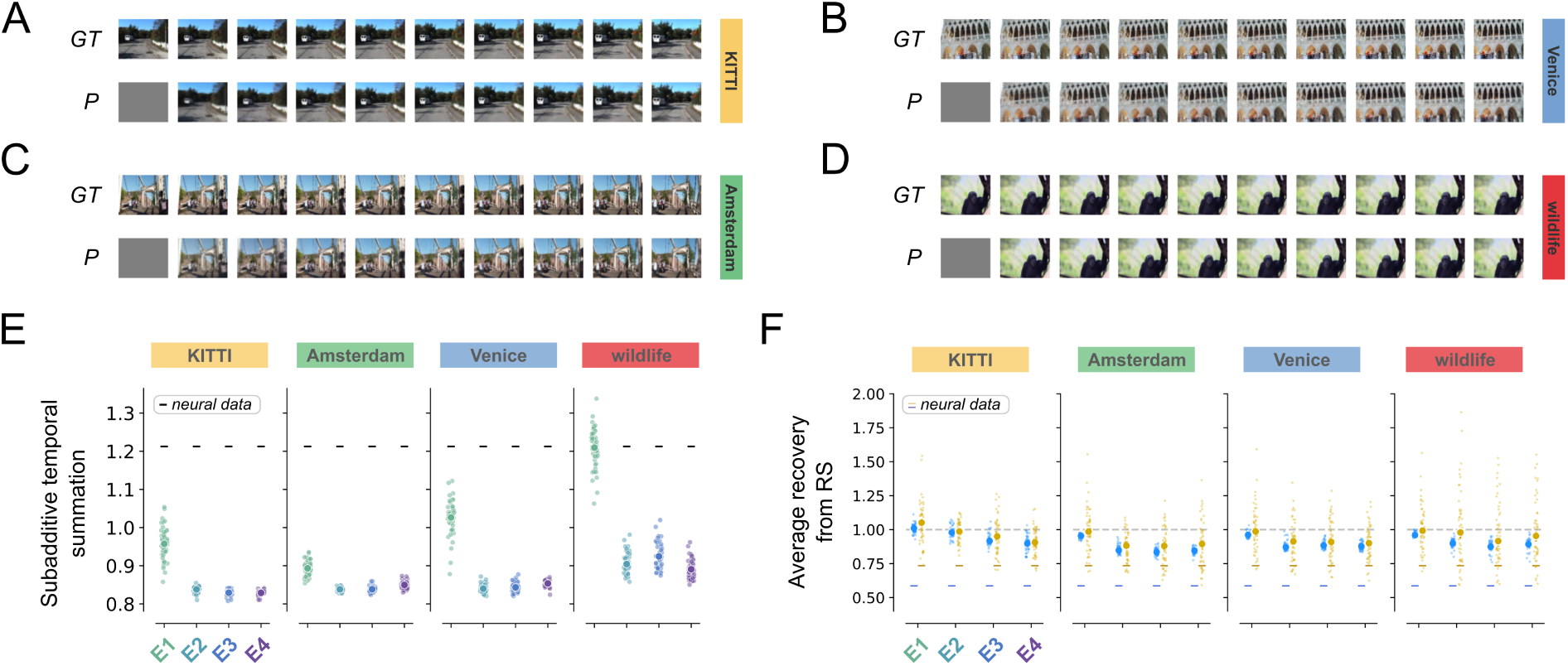
Temporal adaptation signatures in PredNet are robust across datasets. **A-D**: Next frame predictions of PredNet instances trained on four different datasets: the *KITTI* dataset (A) and three video’s from the “Walking Tours” dataset: *Amsterdam* (B), *Venice* (C) and *wildlife* safari (D). **E**: Subadditive temporal summation in PredNet instances trained on the different datasets (panel A-D), determined by computing the ratio between a log and linear fit of the summed responses over varying stimulus durations, for the error units in layer 1 to 4 (E1, E2, E3 and E4, respectively). Data points represent individual images (*n* = 48) and the errorbars depict the SEM across network initializations (*n* = 3). The horizontal lines depict the values derived from the neural data averaged over all visual areas. **F**: Average degree of recovery, determined by computing the response suppression for same (blue) and different (yellow) stimulus pairs averaged over all inter-stimulus intervals, for E1-4. Same network instances and stimuli as depicted in panel (E). The dotted gray line depicts a recovery of 1 (when the magnitude of the first and second response is the same). The horizontal lines depict the values derived from the neural data across all visual areas for same (blue) and different (yellow) stimulus pairs.

In line with our findings in the pretrained PredNet, unit responses of model instances trained on the different datasets all exhibit most pronounced subadditive temporal summation in the first layer (**Fig. 5E**). All the network instances also again show much faster recovery from response suppression compared to the neural data and no repetition suppression (**Fig. 5F**). Notably, while results are thus generally similar across datasets, PredNets trained on the *wildlife* dataset seem to have somewhat distinct dynamics compared to the other three datasets, showing relatively stronger subadditive temporal summation in all layers. We hypothesize that this stronger subadditivity results from the presence of a higher degree of motion continuity in this training dataset, referring to the smooth and natural progression of movement between frames without abrupt transitions, putatively resulting in more accurate predictions (and lower errors) of future frames. Two observations support this hypothesis. First, the *wildlife* dataset shows the highest temporal auto-correlation across video frames across all four datasets (**Supp. Fig. 4B**). Second, we expect that this high auto-correlation results in image sequences that are more similar to static image sequences. Consequently, we expect that networks trained on datasets with higher motion continuity will be less affected if they are trained on static samples from the dataset instead of the full dynamic sequences. Indeed, network instances trained on the *wildlife* dataset show more comparable performance when trained on dynamic versus static sequences compared to the other three datasets (**Supp. Fig. 4C**). These results suggest that training on videos with higher motion continuity leads to lower errors and stronger subadditivity of the unit responses to static stimuli. To conclude, we show that the observed emergent temporal dynamics in PredNet are robust across multiple training datasets, whereby training on more naturalistic, smoothly varying inputs appears to result in more pronounced, visual cortex-like temporal subadditivity of the network responses.

### A weighted loss across layers results in stronger subadditive temporal summation

To examine whether PredNet’s ability to capture temporal adaptation signatures in human visual cortex increases when training with a more biologically plausible loss minimization, we next investigated PredNets that use a layered predictive process by minimizing error across all, instead of solely the first network layer. So far, our PredNets were trained by computing the loss solely over the first network layer (*L*_0_). Next, we trained an additional set of PredNets instances, whereby the loss function additionally receives a weighted sum of errors from the upper network layers (*L_all_*; see Materials and Methods, *Dataset and training manipulations*). The loss of all PredNet instances converged at the end of training, whereby an *L_all_* loss resulted in a slightly better performance compared to an *L*_0_ loss for all datasets (mean*±*SEM, *L*_0_ vs *L_all_*, KITTI: 0.11 vs 0.07; Amsterdam: 0.18 vs 0.12; Venice: 0.11 vs 0.08; wildlife: 0.03 vs 0.02; **Supp. Fig. 5**). We find that the loss indeed influences emergent network dynamics for duration-varying stimuli: PredNet instances trained with *L_all_* exhibit stronger subadditive temporal summation compared to PredNets trained with *L*_0_, for all datasets and across all network layers (**Fig. 6A**). Interestingly, the relative degree of temporal subadditivity in the first layer of PredNets trained with *L_all_* is on par with the neural data (except for the PredNet instances trained on the *wildlife* dataset which now exhibit even stronger compression). For the repetition trials, we observe no effect of the chosen loss function on the emergent temporal dynamics (**Fig. 6B**), regardless which dataset was used for optimization, which is most likely due to the fast dynamics we observed in the pretrained PredNet, resulting in rapid recovery of the unit responses and lack of repetition suppression. All together, these results show that optimizing the PredNet by aggregating the loss over all hierarchical layers results in stronger subadditive temporal summation, thereby aligning more closely with the neural data, but does not improve the alignment with visual cortex responses to repeated stimulus presentations.

**Figure 6:**
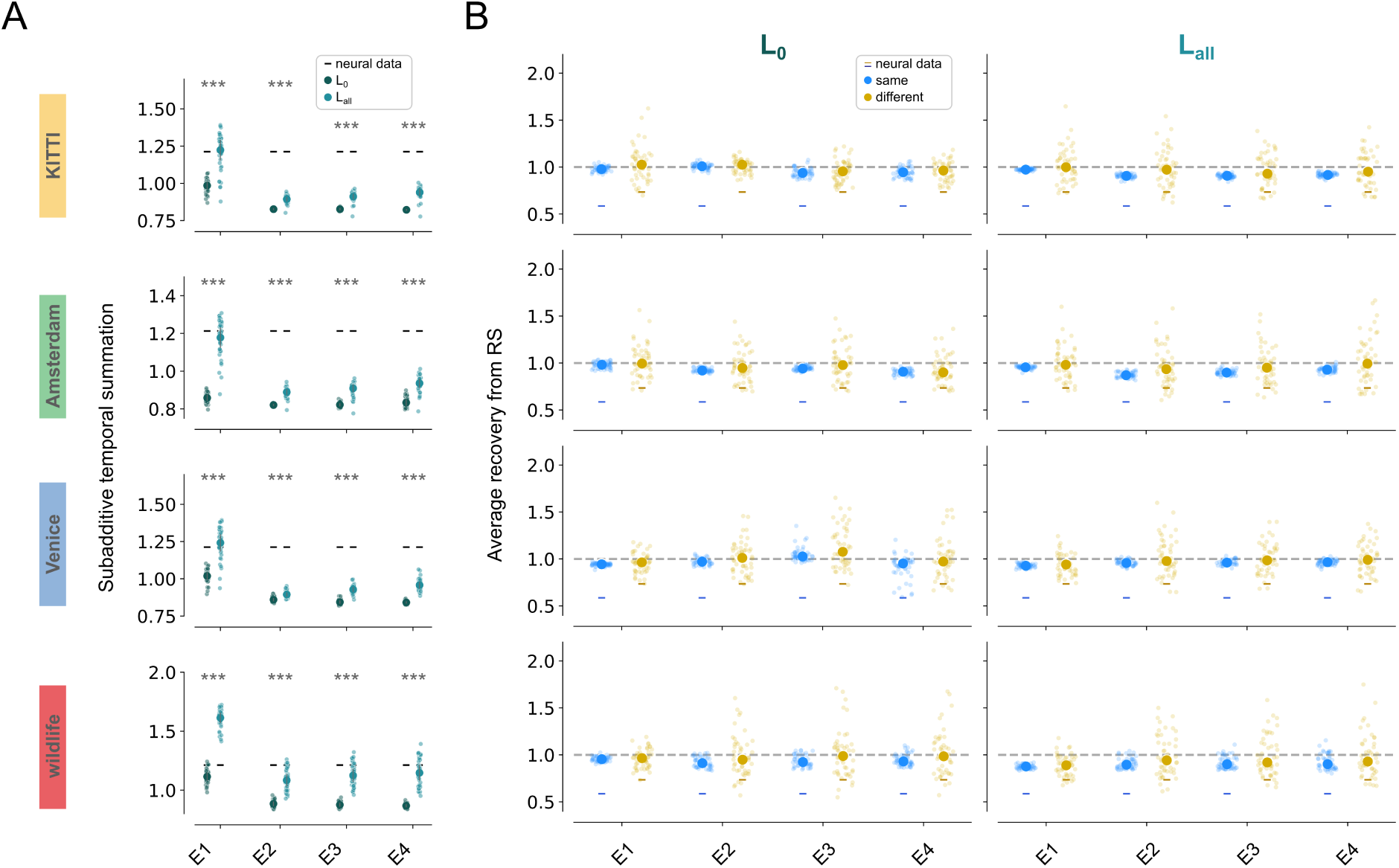
Error-minimization across all layers increases subadditive temporal summation. **A**: The degree of subadditive temporal summation, determined by computing the ratio between a log and linear fit of the summed responses over varying stimulus durations, for the error units in layer 1 to 4 (E1, E2, E3 and E4, respectively). PredNet instances (*n* = 3) were trained on one of the four datasets (rows) with an *L*_0_ (green) or *L_all_* (blue) loss. Data points represent individual stimuli (*n* = 48) averaged over network instances and the error bars depict the SEM across the stimuli. The horizontal bars depict the values derived from the neural data averaged over all visual areas. Independent T-test, ^∗∗∗^ p *<* 0.001. **B**: Average degree of recovery, determined by computing the response suppression for same (blue) and different (yellow) repeated stimuli averaged over the inter-stimulus intervals, for the error units in layer 1 to 4. Same network instances and stimuli as depicted in panel (A). The horizontal bars depict the values derived from the neural data across all visual areas for same (blue) and different (yellow) repeated stimulus presentations, respectively.

### Sustained level activity originates from cluttered parts in the image

So far, we investigated whether the PredNet captures neural signatures of temporal adaptation in human visual cortex. While PredNet surprisingly lacks repetition suppression effects, it exhibits similar temporal response summation and the associated transient-sustained dynamics as observed in human visual cortex. Moreover, we found that the degree of temporal summation in the network is modulated by motion continuity and the loss computation, with more biologically plausible training regimes resulting in more cortex-like temporal summation. But how does this subadditive summation relate to PredNet’s ability to predict the next frame? In the last two sections, we uncover a relation between subadditivity and spatial image properties by visualizing and quantifying error unit activations over time for individual image pixels.

Visualization of error unit activations in the pretrained PredNet for an example image shows that the decay to sustained response level for longer stimulus durations is caused by the network generating increasingly accurate predictions of the visual scene (**Fig. 7A**). Notably, error unit activity seems to be mostly driven by the parts of the image that contain more contours and edges. This spatially localized increase in activations is most pronounced for the first layer, whereas later layers show more distributed activations over space, likely due to the max-pooling operation applied on the representation units. To illustrate the influence of image clutter on sustained error unit activity, we directly compare PredNet responses for two different example images: a relatively simple scene containing several homogeneous sections along with a few contours (**Fig. 6B**, *top*) versus a more complex scene with many small objects, thus containing many contours and a high degree of clutter (**Fig. 7B**, *bottom*). Since error activity is localized to more cluttered parts of the scenes, the bottom image that has more clutter shows on average a higher sustained response (**Fig. 7C**). Consistent with these observations in the pretrained network, we find that the stronger subadditive temporal summation for PredNets trained in-house with an *L_all_* loss was associated with better predictions in cluttered parts of the image (**Supp. Fig. 6**). Together, these results suggest that the sustained level activity is predominantly caused by the network’s lack of precise predictions for the cluttered image locations.

**Figure 7:**
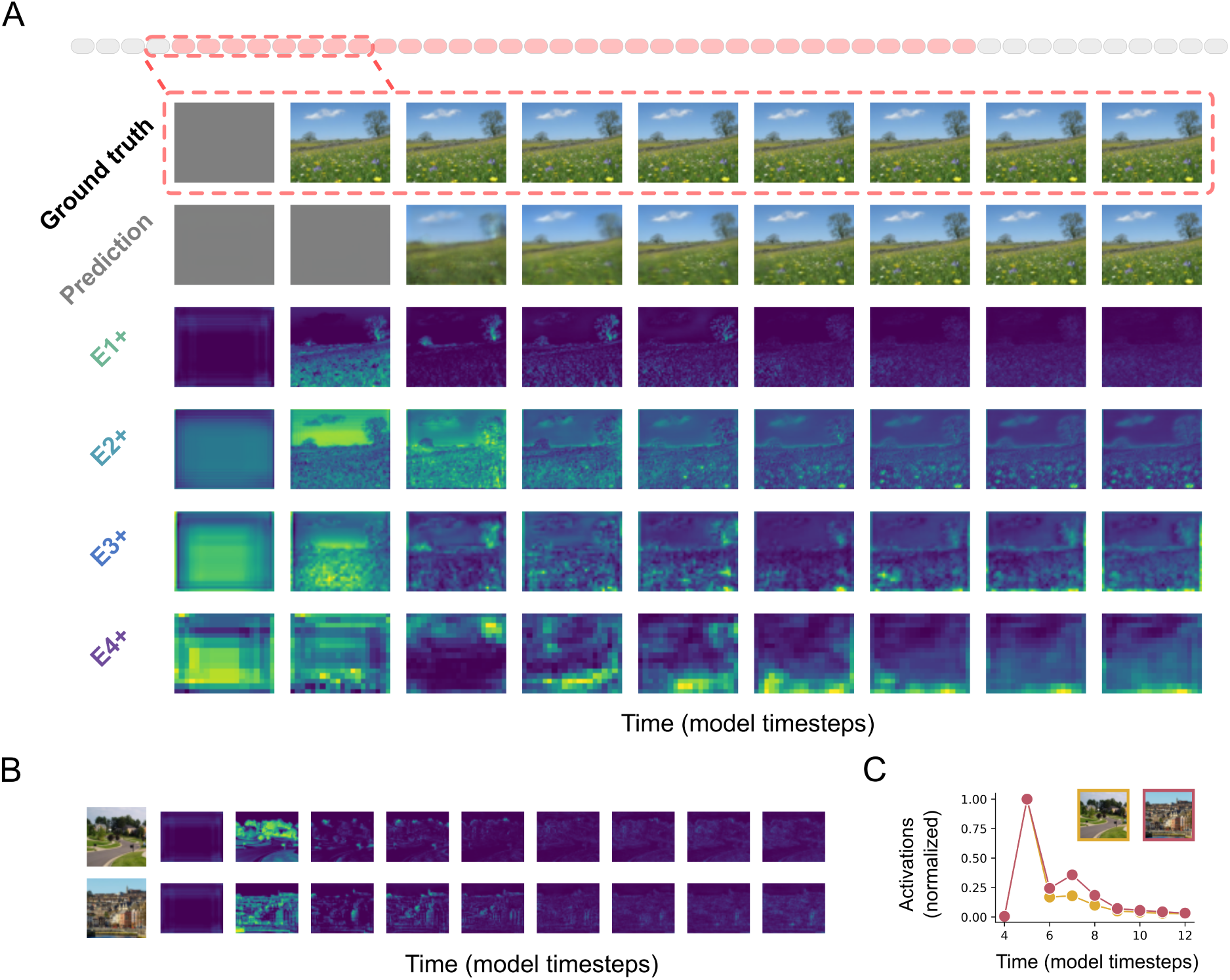
Activity in error units originate from cluttered parts of the image. **A**: PredNet activations for a single stimulus trial for an example image. The top two rows depict the presented image (Ground truth) and the output of the PredNet (Prediction) at timestep *t*. The four rows below depict the activations of positive error units shown separately for each of the four layers. Higher error activity is indicated in brighter colors. At the first image representation (*t*_4_) the activation of error units increases in all layers. For the successive timesteps, the PredNet adjust its prediction resulting in a decrease in the activation of the error units in all layers, converging to a sustained level caused by edges and contours. **B**: Activity of error units for two images, consisting either of a sparse scene with relatively few edges and contours (*top*) versus a more cluttered scene (*bottom*). **C**: Normalized activations of the error units depicted for the two images depicted in panel (B). The more complex scene gives rise to higher sustained level activity of the error units compared to the sparse scene.

### Degree of sustained-level activity in PredNet is linked to low-level scene statistics

To investigate the relationship between error responses and image clutter more systematically, we correlated the average sustained level activity of error units for each image with two biologically plausible natural scene statistics which capture the degree of clutter in an image, contrast energy (CE) and spatial coherence (SC) (see Materials and Methods, *Summary metrics*). Cluttered or complex scenes tend to have higher CE and CS values, while sparse scenes have low values (**Fig. 8A**; Scholte et al. 2009; Groen et al. 2013). We indeed find a significant correlation between sustained PredNet error unit activity and the two scene statistics in the first network layer (**Fig. 8B**): images with high CE and SC values give rise to increased sustained response magnitude relative to the peak, evident by low transient-to-sustained ratios (linear regression, CE, *R*^2^ = 0.22, *p* = 0.001; SC, *R*^2^ = 0.13, *p* = 0.012), whereas images with low CE and SC values show a reduced sustained response magnitude relative to the peak, associated with high transient-to-sustained ratios.

**Figure 8:**
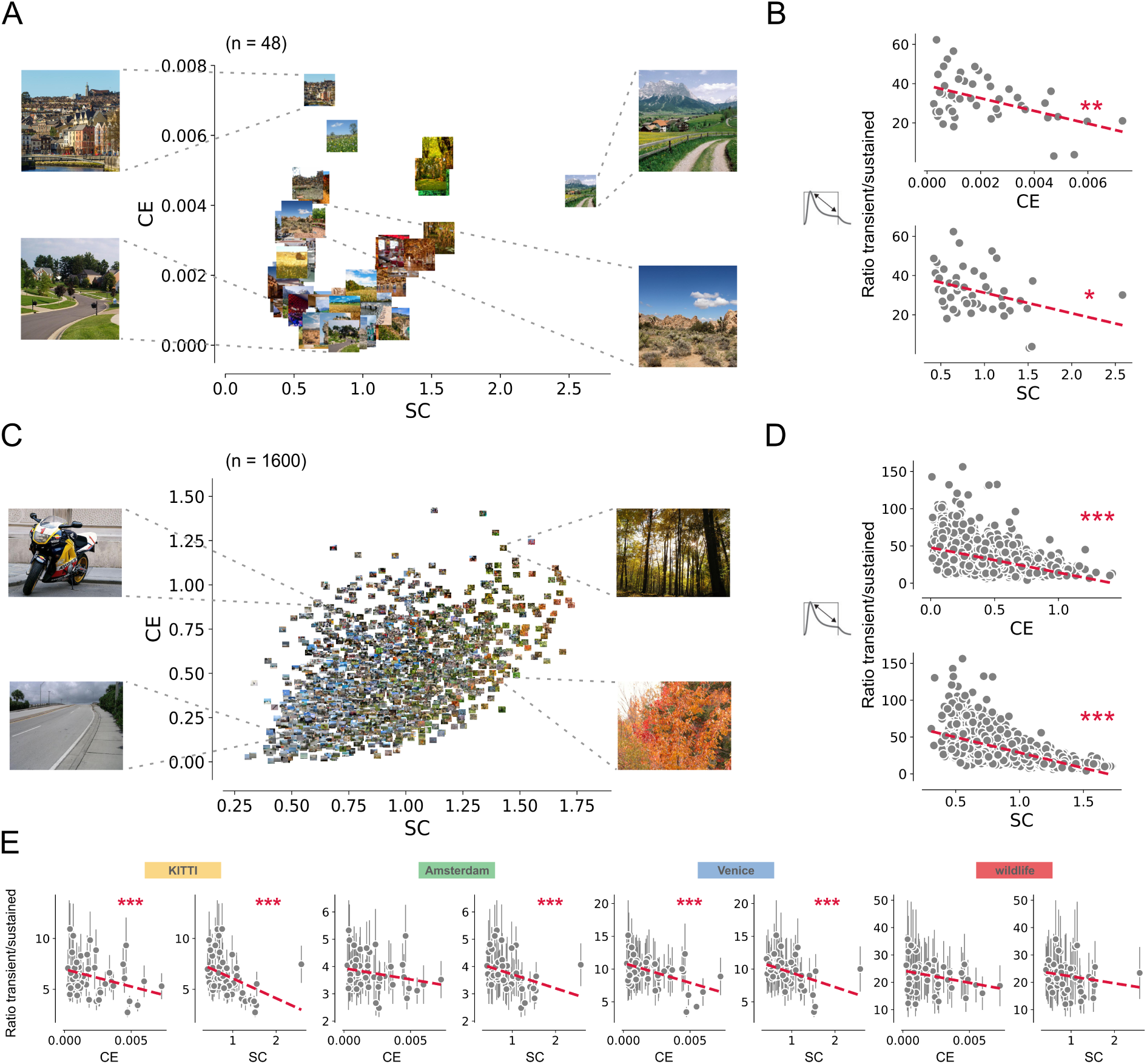
Higher sustained activity for images depicting cluttered scenes. **A**: A subset (n = 48) of the images belonging to the dataset curated by Brands et al. (2024) plotted against CE and SC values. CE represents the approximation of the *β* parameter of the Weibull function and varies with the distribution of the local contrasts strength. SC is the approximation of the *γ* of the Weibull function and varies with the amount of scene fragmentation or scene clutter. **B**: Ratio between the transient and sustained response of the error units belonging to the first layer of the PredNet against the CE (*top*) and SC (*bottom*) values of the images depicted in panel A. **C-D**: Same as A-B for the images (n = 1600) belonging to the dataset curated by Groen et al. (2013). **E**: Ratio between the transient and sustained response of the error units belonging to the first layer of the PredNet against the CE (*top*) and SC (*bottom*) values for the different datasets. Dots represent individual images and the errorbars depict the SEM across network initializations. ^∗∗∗^ p *<* 0.001.

To assess the robustness of this correlation, we repeated this analysis using a larger, independent stimulus dataset that covers a broader range of CE and SC values (**Fig. 8C**), which uncovered the same association in both the pretrained PredNet (transient-sustained ratio vs. CE, *R*^2^ = 0.21, *p <* 0.001; transient-sustained ratio vs. SC, *R*^2^ = 0.39, *p <* 0.001; **Fig. 8D**) and three of the four PredNets we trained ourselves, namely the *KITTI* (CE, *R*^2^ = 0.09, *p* = 0.039; SC, *R*^2^ = 0.17, *p* = 0.004), *Amsterdam* (SC, *R*^2^ = 0.115, *p* = 0.019) and *Venice* (CE, *R*^2^ = 0.23, *p* = 0.001; SC, *R*^2^ = 0.168, *p* = 0.004; **Fig. 8E**) dataset. However, PredNets trained on the *wildlife* dataset did not show a significant correlation (CE, *R*^2^ = 0.08, *p* = 0.051; SC, *R*^2^ = 0.04, *p* = 0.2). This may be due to the overall higher range of CE and SC values in the video frames of this dataset (**Supp. Fig. 4D**), allowing the network to generates good predictions for these images and hence less prolonged error activations. Importantly, the significant relation between image clutter and transient-sustained dynamics is exclusive to the first network layer; layers *E2*-*E4* do not show this (**Supp. Fig. 7**), possibly due to the more distributed activations over space for these network layers (**Fig. 7A**). These results suggest that degree of sustained error activity - and thereby the degree of temporal response adaptation - on a given trial in PredNet is related to the complexity of both the presented test image as well as the images it was trained on.

Overall, these correlations between temporal dynamics of first-layer PredNet units and image complexity suggest that the observed brain-like signatures of short-term temporal adaptation in PredNet are mostly tied to low-level properties of the visual inputs. While our prior work has shown that image complexity modulates EEG activity at distinct temporal stages of processing (Groen et al., 2013), and we have proposed that higher complexity images may reflect the need for more extended recurrent processing (Groen et al., 2018), we have not directly linked these low-level properties to short-term adaptation dynamics. To determine whether sustained responses in our intracranial recordings were also enhanced for more cluttered images, we performed the same analyses on our iEEG broadband responses. We indeed find a trend of higher transient-to-sustained ratio for lower SC values (linear regression, *R*^2^ = 0.34, p = 0.06, **Supp. Fig. 8**). While the small number of test images here precludes us from drawing firm conclusions, these results hint at the possibility that transient-sustained dynamics in human visual cortex and the resulting degree of temporal subadditivity are also shaped by low-level image properties.

## Discussion

The aim of this study was to perform a critical test of the biological fidelity of predictive coding as a theory of cortical function. To this end, we investigated and compared emergent temporal dynamics exhibited by a predictive coding network, PredNet, with the temporal dynamics observed in a neural dataset of iEEG broadband responses when presented with single or repeated stimulus presentations. For single stimulus presentations, we demonstrate that error unit activations in the first layer of the PredNet exhibit subadditive temporal summation, but, as opposed to the neural data, also show systematic offset responses. For repeated stimulus presentations, error unit responses show a slight reduction in response magnitude when an image is preceded by another, but fail to show repetition suppression as observed in the neural responses, which is considered a robust and ubiquitous phenomenon across the brain. Computing the loss over all layers, as opposed to solely over the first, increases the degree of subadditive temporal summation, thereby showing stronger overlap with the neural data, but does not affect the repetition suppression patterns. Lastly, we reveal that complex images containing a high degree of clutter lead to higher levels of sustained responses and reveal a similar trend in the iEEG broadband broadband responses. These findings show that the PredNet’s emergent temporal dynamics only partly capture temporal adaptation signatures of human visual cortex, thereby suggesting that predictive coding, as instantiated in PredNet, does not fully account for cortical dynamics observed in the brain.

### Nonlinear dynamics in human visual cortex and predictive coding networks

In human visual cortex, neural responses show adaptation over time, thereby exhibiting subadditive temporal summation and repetition suppression to single and repeated stimuli, respectively (Zhou et al., 2019; Groen et al., 2022; Brands et al., 2024). Here, we confirm these two signatures of neural adaptation in our iEEG dataset, evident by the fact that neural responses in both early and late visual areas exhibit subadditivity of the response magnitudes with prolonged stimulus durations and stronger response suppression for same as opposed to different inputs when shown in sequence. We show that PredNets capture some of the dynamics present in the neural data, namely subadditive temporal summation, but fails to reproduce others, namely repetition suppression. To determine the fidelity of the PredNet to biological systems, it is helpful to ask which features and components of the model are responsible or necessary to reproduce these neural dynamics. For example, one key feature of the PredNet is recurrent connectivity, which is required to observe temporal dynamics, since a strictly feedforward version of the model would lack temporal dynamics all together. Another important PredNet feature, motivated by predictive coding principles (Rao and Ballard, 1999), is that the network explicitly computes an error representation in a population of neurons, which is propagated from layer to layer in a feedforward manner and is used to update the network representations and consequently, network predictions. These two PredNet features - recurrence and explicit error representation - presumably allow the network to exhibit several of the neural response properties observed in the neural data. For duration trials, these network features result in subadditive temporal dynamics, characterized by transient-sustained dynamics, with an initial prediction error at stimulus onset, and a decay of the error as a consequence of the model updating its predictions. For repeated stimuli, these network features result in a slightly lower error during the second compared to first stimulus presentation, especially for when two stimuli are in close temporal proximity of each other and the two stimuli are the same, which is the result of the lingering prediction of the first stimulus presentation during the presentation of the second. However, due to the fast updating of the network representations, the network “forgets” previous inputs as the time in between stimuli increases, resulting in a full recovery of the response suppression for short ISIs and no difference between same and different stimuli, thereby deviating from our observations in the neural data. These results demonstrate that while features as recurrence and explicit error representation may effectively capture some of the neural signatures of temporal adaptation, there is a misalignment between “biological” time and the notion of “time” in the PredNet. Additional features, either on the side of the inputs (e.g. adjusting the sample rate of the frames) or in PredNet itself (e.g. controlling the rate with which representations are updated) might be necessary to flexibility adjust the temporal resolution of the PredNet such that it better matches that of the neural data.

### Discrepancy between human iEEG and PredNet in offset responses

One discrepancy we find between PredNet unit activation timecourses and broadband iEEG responses is the prominent offset response exhibited by the PredNet and the lack of this offset response in the neural data. In the PredNet, the strong offset response is a direct consequence of the explicit representation of the error activations and a training objective based on reducing these errors, resulting in a large prediction error and second activation peak to the offset of the stimulus. In our data however, we did not observe strong stimulus offset responses. Previous studies have shown inconsistencies regarding the presence of offset dynamics in neural responses, with some iEEG dataset showing a similar lack (Zhou et al., 2019; Groen et al., 2012a), while other human iEEG (Zhou et al., 2019) and animal (Bair et al., 2002; Benucci et al., 2009) datasets did show a response offset in at least a subset of neural responses. From a methodological perspective, the presence of an offset response in the neural timecourses can depend on several factors, such as the data type used for data collection (e.g., fMRI vs. iEEG), brain areas sampled, or experimental design. From an empirical perspective, it is still debated what causes the presence of offset responses or the lack thereof. A previous study noted that offset responses in an iEEG data were more pronounced for electrodes with peripherally tuned spatial receptive fields, suggesting a link between the offset response and spatial coverage of the stimulus (Zhou et al., 2019). Other work has hypothesized offset dynamics are neural representations which reflects differences in information processing, whereby transient responses both on the on- and offset of stimuli indicate involvement in detecting temporal change, while the absence of an offset is related to other types of information processing, including object recognition and appearance (Zhou et al., 2018). Moreover, earlier work has proposed segregated neural pathways for onset and offset responses as a feature of many sensory computations, for example in motion detection (Westheimer, 2007), retinal information processing (Westheimer, 2007), perceptual grouping of auditory stimuli (Bregman, 1994) and olfaction-related behavior (Chalasani et al., 2007). Our findings suggest that predictive coding theory alone may not be sufficient to fully capture the heterogeneous offset response profiles observed for single, duration-varying stimuli in neural data; additional features, such as spatial topography or separate pathways for predicting motion and object identity, may be needed to better capture the full range of neural responses.

### The role of error minimization on temporal adaptation

While previous work has studied the effect of layer-specific error minimization (Lotter et al., 2016; Rane et al., 2020), these studies have focused on network performance and did not compare the resulting emergent temporal dynamics with the dynamics observed in neural data. Aside from our empirical result which suggests multi-layer error minimization results in dynamics more closely mimicking that of biological systems, it is also a more plausible learning objective given how the brain processes sensory information. More specifically, the abundant presence of feedback connections in the visual cortex (Johnson and Burkhalter, 1996; Lamme et al., 1998) in combination with the influence of these feedback connections on synaptic plasticity (Tropea et al., 1999; Roelfsema and Holtmaat, 2018) implies that the brain’s learning rules are applied across multiple hierarchical layers, thereby shaping the functional structure (and consequently dynamics) of multilayered neuronal networks. Another argument for computing the error over all layers is the prevention of emergent sub-modules, whereby Rane et al. (2020) showed that when PredNet is trained solely on the first layer, the lowest layer focuses on realistic predictions, while higher layers act as a single network regressing onto the first layer’s units. This prevents higher layers from developing distinct predictive dynamics, contradicting the hierarchical structure of the visual cortex, where each area has unique dynamics and extracts different features from visual inputs. Future directions could further elucidate the effect of layer-specific error minimization, by for instance varying the weights for aggregating the error across layers and investigate how this affects the increase in subadditivity exhibited by the unit responses. Another direction could be to not aggregate the error across layers, but instead perform layer-specific updates of the network during training, such that more emphasis is imposed on accurate higher-level representations. These changes to the loss computation may help gain better insights into how the explicit error representations across layers during optimization affects emergent temporal dynamics in the network, and which objectives yield dynamics which more closely resemble that of biological systems.

### PredNet reveals a link between sustained responses and low-level image statistics

We find a relationship between the degree of clutter in an image and the sustained response magnitude of PredNet error units when presented with single images presented for longer durations. Moreover, we observe a similar trend in the iEEG broadband responses, whereby sustained level responses are higher for more cluttered images. This is in line with previous studies investigating the effect of image statistics in evoked neural responses using EEG (Scholte et al., 2009; Groen et al., 2013; Seijdel et al., 2020), whereby results showed that both CE and SC modulate response amplitudes over time (Groen et al., 2013). We have previously suggested that extracting these statistical properties of the visual input enables accurate and efficient perception, for example for judgments of texture invariance (Groen et al., 2012a) and visual similarity (Groen et al., 2012b). Moreover, fMRI studies showed that both low- and high-level brain regions are sensitive to the degree of clutter in a scene (Park et al., 2015) and that scene complexity modulates the degree of feedback activity during object detection in natural scenes (Groen et al., 2018).

In the PredNet, the transition from transient to sustained level activity of the unit activations is a direct result of improved predictions made by the network. Therefore, one interpretation of this relation between error unit activity and image complexity in PredNet is that the degree of clutter in an image directly affects the predictivity of the scene. Specifically, as the complexity of a scene increases, the brain’s predictions may become less accurate, requiring more top-down feedback to adjust and refine sensory processing. This aligns with the idea that feedback signals are crucial for correcting prediction errors (Watabe-Uchida et al., 2017), particularly in more uncertain or complex environments. However, we have previously shown that short-term adaptation phenomena in intracranial EEG timecourses, including transient-sustained dynamics (Zhou et al., 2019; Groen et al., 2022; Brands et al., 2024), are well captured by divisive normalization mechanisms, which compute the time-varying neural response in a bottom-up manner from the ratio between a linear response timecourse divided by the summed activity of a larger pool of neurons (Heeger, 1992, 1993; Carandini and Heeger, 2012). Unlike the PredNet, these mechanism accurately predict repetition suppression in the neural responses, demonstrating that it is possible to capture temporal adaptation phenomena in human visual cortex without higher-level prediction mechanisms.

### Limitations and future work

First, since neural recordings were collected from few subjects presented with a small number of images, interpretation regarding the relationship between the degree of clutter in the image and the subadditivity of neural responses should be made with caution. Analyzing additional datasets will help gain insights whether the predictions made by the PredNet regarding the relationship between low-level statistical regularities and the response magnitude of the sustained level align with biological systems. Second, in the current study, we focused on the temporal dynamics of the error units. However, to determine whether predictive coding is a faithful model of cortical function, other computations made within the network (e.g. by the representation units) should be studied and compared to neural data as well. Lastly, there has been some debate about whether the PredNet fully embodies the principles of predictive coding (Rane et al., 2020). To determine whether predictive coding accounts for short-term temporal adaptation in human visual cortex, future research should explore alternative implementations to verify that the our findings are not specific to the current architecture, but generalize to other implementations (e.g. Heilbron and de Lange 2023) of the predictive coding scheme as well. All together, these findings highlight the potential of PredNet in modeling certain aspects of temporal adaptation, while also showing misalignments with the neural data, suggesting that predictive, top-down processes are not sufficient, or even necessary, to fully capture the richness of temporal dynamics in human visual cortex.

## Supplementary Tables

**Supplementary Table 1:** Overview of video datasets used to train the PredNet. Columns refer to the following: Dataset, name of the dataset. Duration: length of the video, HH:MM:SS. No. frames, Number of frames in the video. Fps, framer-per-second. HxWxC, Input of the input, height×width×. Mounting, View of the camera, either mounted on a car of a person.

| Dataset | Duration | No. frames | fps | input size (H × W × C) | Mounting |
| --- | --- | --- | --- | --- | --- |
| KITTI | 00:34:30 | 41.396 | 15 | 128 × 160 × 3 | car |
| Walking Tours - Amsterdam | 00:30:33 | 27.500 | 15 | 128 × 160 × 3 | person |
| Walking Tours - Venice | 01:01:07 | 55.000 | 15 | 128 × 160 × 3 | person |
| Walking Tours - wildlife safari | 01:00:04 | 54.058 | 15 | 128 × 160 × 3 | person |

## Supplementary Figures

**Supplementary Figure 1:**
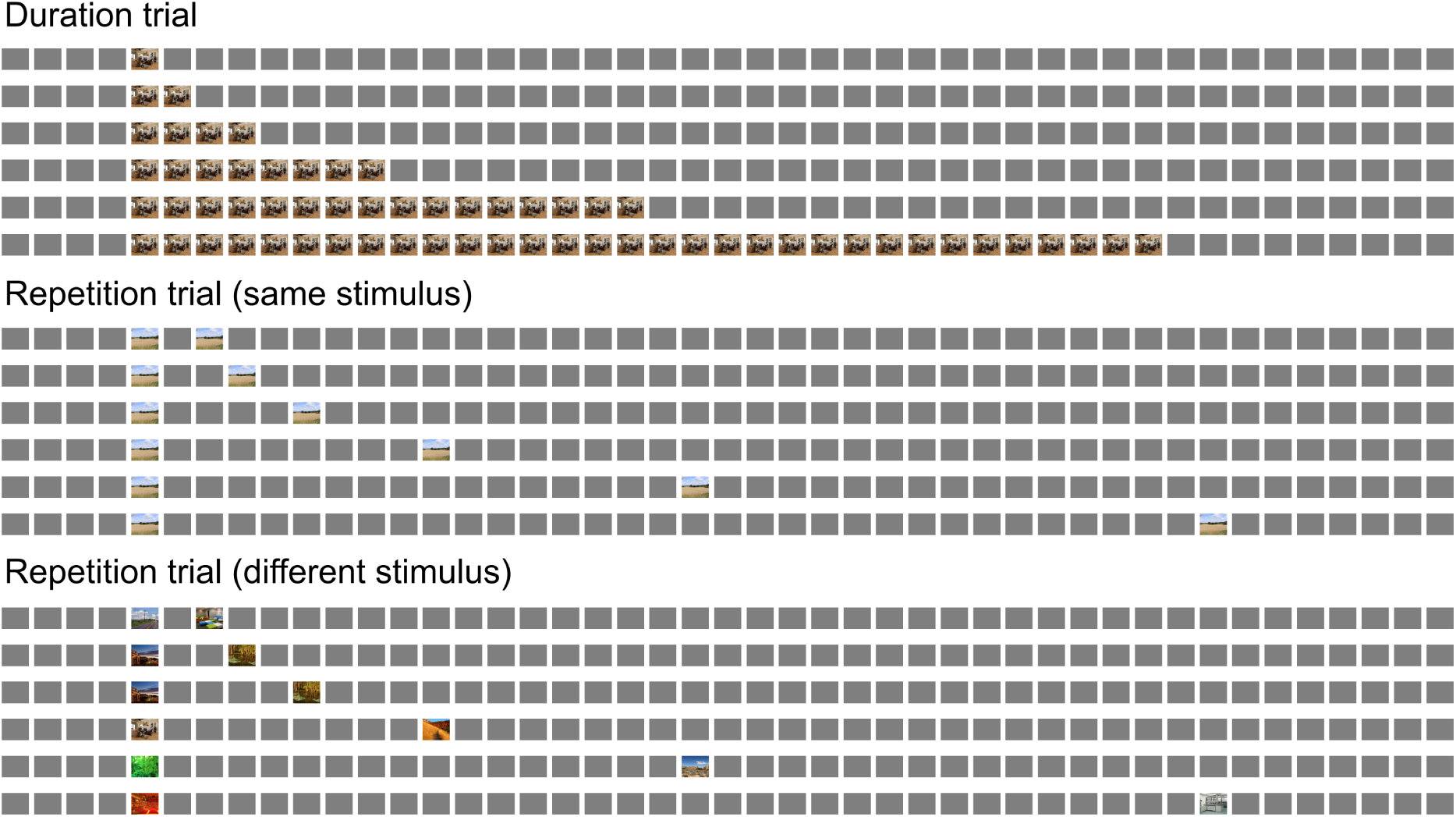
Example sequences of stimuli presented to the PredNet. Duration trials (top) consisted of presentations of a single image for a varying duration (total sequence length of 45 timesteps), defined in powers of two, i.e. 1, 2, 4, 8, 16 and 32. Repetition trials consisted of the presentation of two images which are either the same (middle) or different (bottom) with varying inter-stimulus intervals (same temporal step size as the single stimulus trials, i.e. 1, 2, 4, 8, 16 and 32).

**Supplementary Figure 2:**
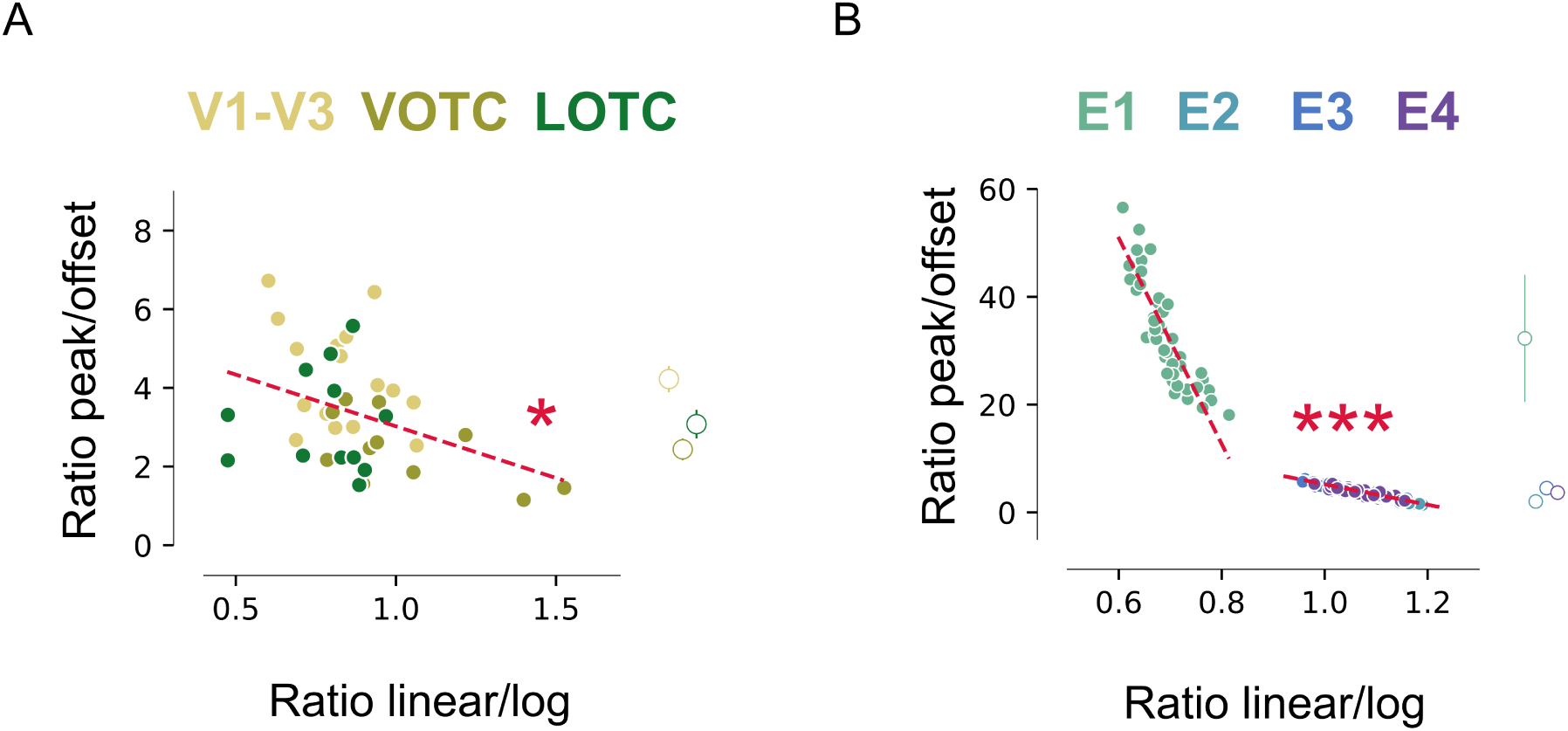
Correlation between subadditive temporal summation and transient-sustained response dynamics. **A**: The ratio between the transient and sustained response against the ratio of the explained variance by a linear and logarithmic curve fit for the summed response magnitude over the different stimulus durations for V1-V3, ventral occipito-temporal cortex (VOTC) and lateral occipito-temporal cortex (LOTC). Each solid marker depicts an individual electrode (V1-V3, *n* = 17; VOTC, *n* = 11; LOTC, *n* = 13). The area-specific average for the ratio of the transient and sustained response is depicted on the right. **B**: Same as panel (A) for the error units of the PredNet depicted separately for each layer (i.e. E1, E2, E3 and E4). Data points depict individual images (*n* = 48). Linear regression, ^∗^ p *<* 0.05, ^∗∗∗^ p *<* 0.001.

**Supplementary Figure 3:**
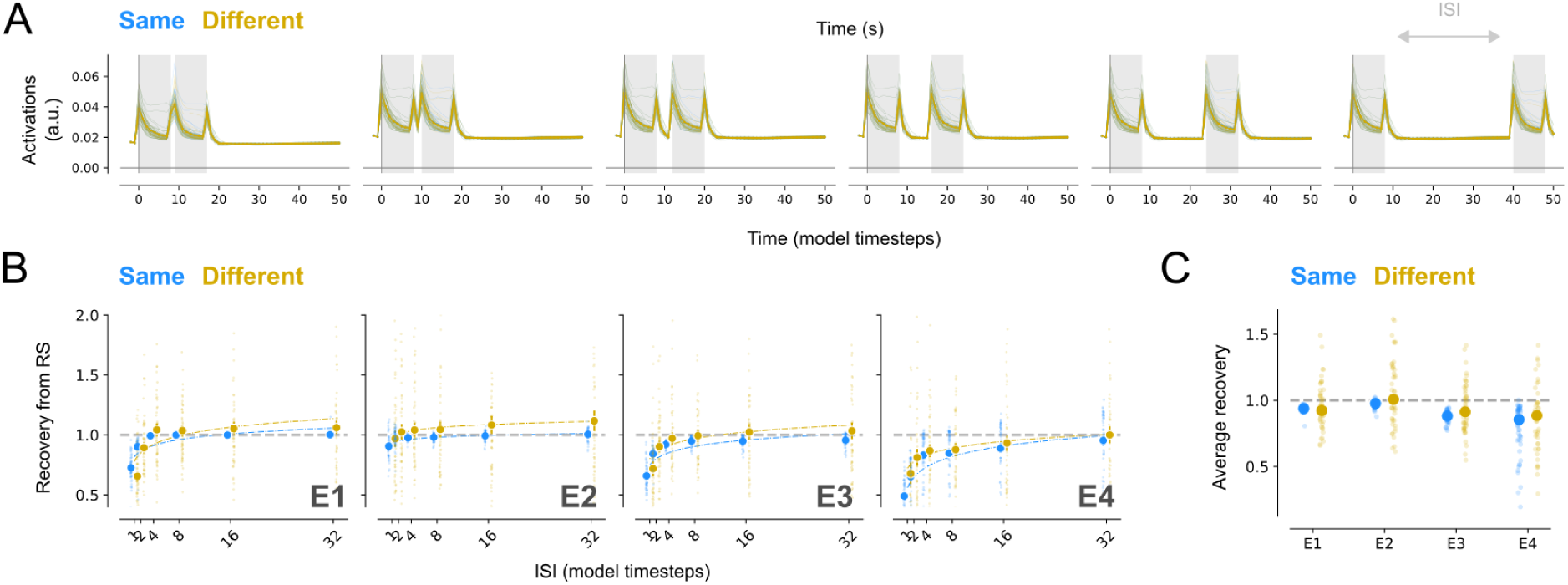
Recovery form repetition for trials with long stimulus durations. **A**: PredNet activations of the error units in the first network layer for the six different interstimulus intervals (ISI) between two stimuli with a duration of 8 model timesteps. **B**: Recovery from response suppression for same and different stimuli plotted separately per network layer (i.e. E1, E2 E3 and E4). Values are obtained by computing the AUC ratios between the first and second stimulus. The fitted curves express recovery as a function of the ISI. The dotted grey line depicts a recovery of 1 (i.e. when the magnitude of the first and second response is the same). **C**: Average degree of recovery computed over all ISIs.

**Supplementary Figure 4:**
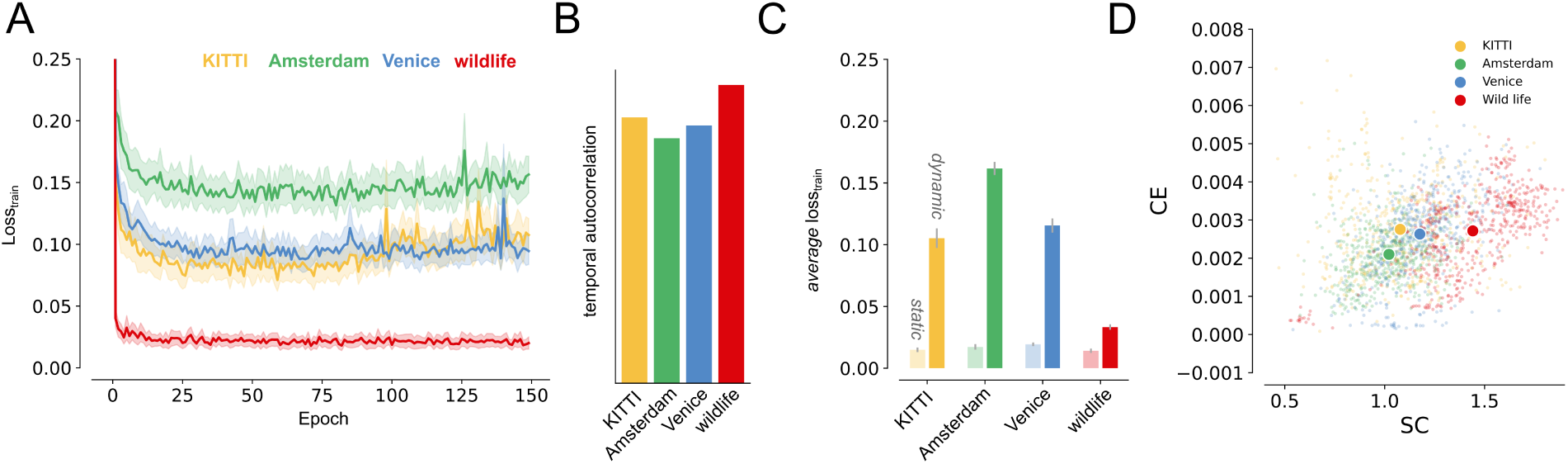
Training loss and image statistics for the different video datasets. **A**: Average training loss across PredNet instances (*n* = 3) trained on the *KITTI* video dataset or one of three video’s from the “Walking Tours” dataset, including *Amsterdam*, *Venice* and from a *wildlife* safari. Curves are smoothed with a Gaussian kernel with standard deviation of *σ* = 10. The shaded region depicts the SEM across network initializations. **B**: Temporal autocorrelation, described by the Pearson product-moment correlation coefficients across video frames. Higher temporal autocorrelation occurs in videos with slow-moving objects or static scenes, where consecutive frames are very similar. Low temporal autocorrelation occurs in videos with fast motion or abrupt changes, where frames differ significantly over short intervals. **C**: Average loss across PredNet instances (*n* = 3) on the four datasets introduced in panel (A), with either static (light) or dynamic (dark) frame sequences. **D**: CE and SC values for a subset of the images (*n* = 2000) randomly sampled from each video dataset.

**Supplementary Figure 5:**
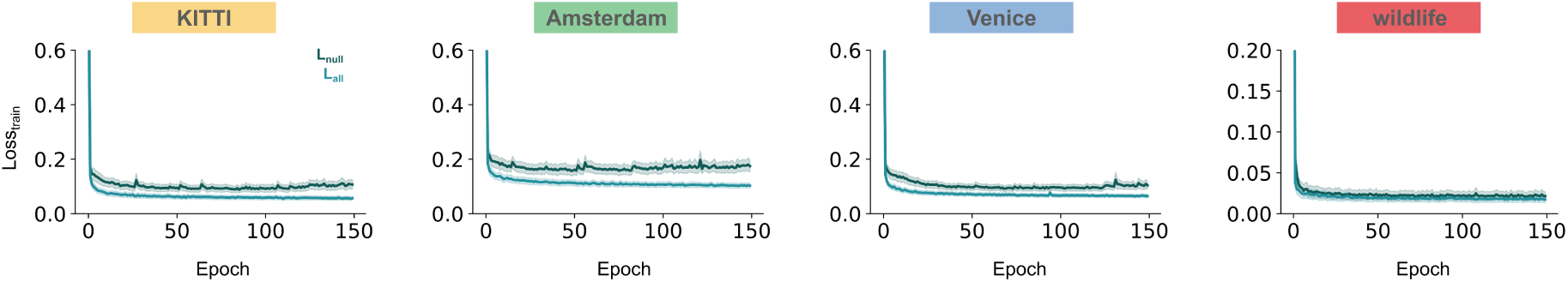
Training loss across video datasets for different objective functions. Average training loss across PredNet instances (*n* = 3) trained with a *L*_0_ or *L_all_* loss on the *KITTI* video dataset or one of three video’s from the “Walking Tours” dataset, including *Amsterdam*, *Venice* and from a *wildlife* safari. Curves are smoothed with a Gaussian kernel with standard deviation of *σ* = 10. The shaded region depicts the SEM across network initializations.

**Supplementary Figure 6:**
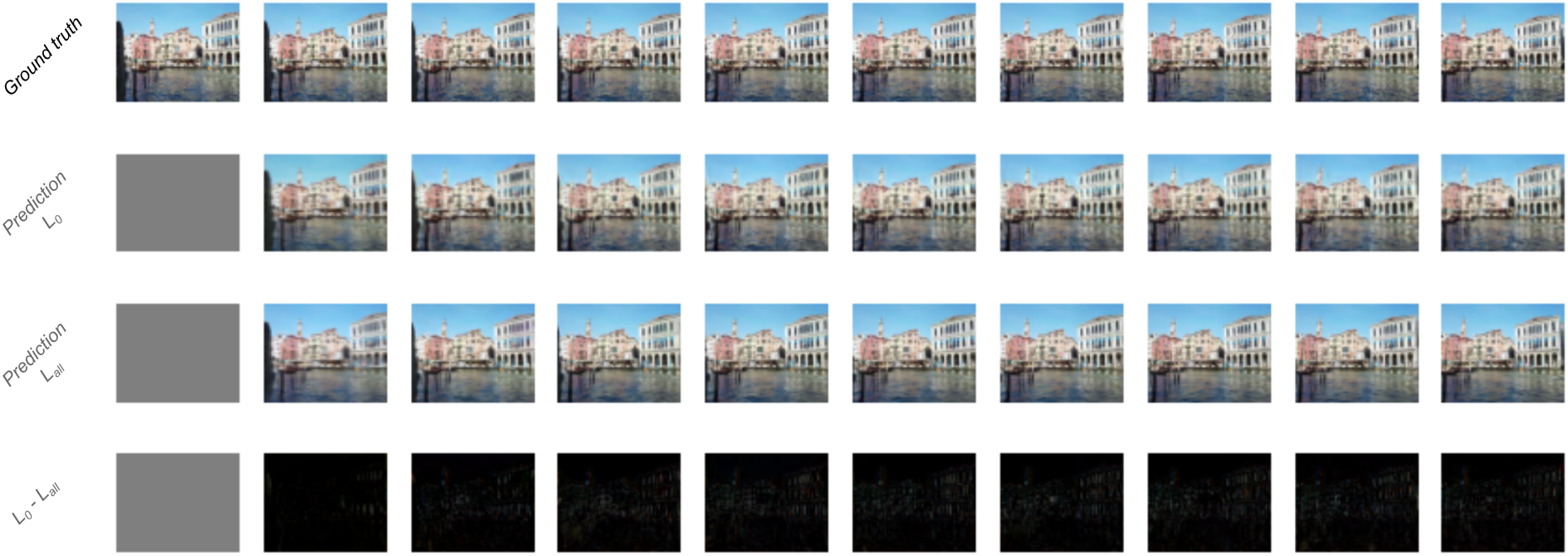
Optimizing the PredNet by computing errors over all layers improves predictions for cluttered parts of the image. Next-frame predictions of PredNet *L*_0_ (first row) and *L_all_* (second row). The “*L*_0_ − L_all_” visualization shows where the predictions differ between models.

**Supplementary Figure 7:**
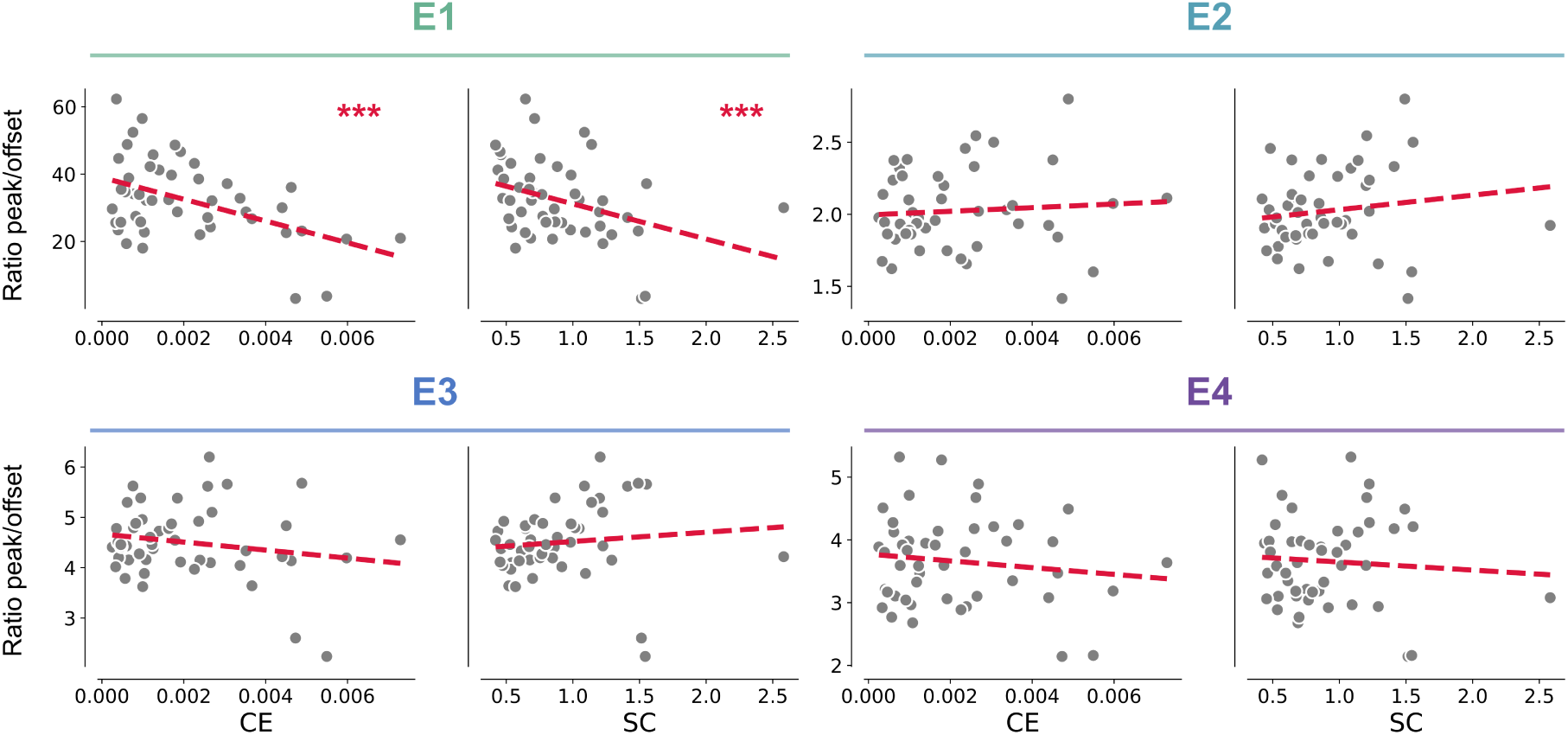
Nonlinearity in temporal dynamics of transient-sustained responses affected by low-level image statistics. Ratio between the transient and sustained response of the error units of the PredNet against the CE (*top*) and SC (*bottom*) plotted separately per layer. The data for for layer 1 (E1) is also depicted in the main text (Fig. 7B). Dots represent individual images. ^∗∗∗^ p *<* 0.001.

**Supplementary Figure 8:**
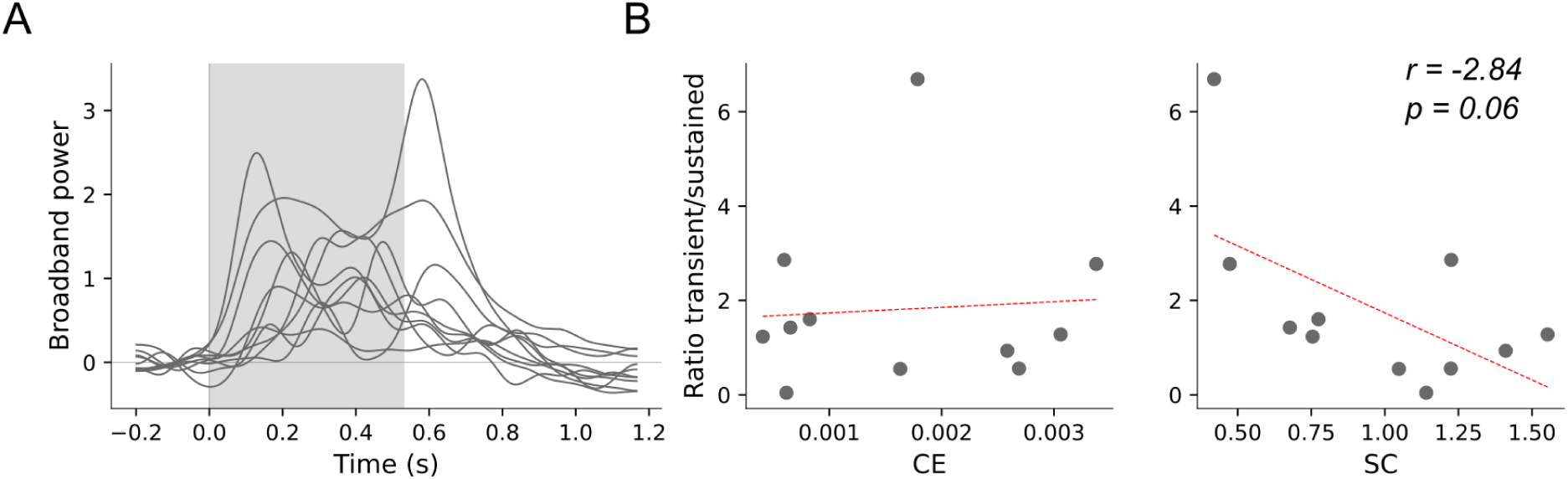
Relationship between sustained level activity and low-level image statistics in iEEG broadband data. **A**: Broadband responses for individual images for a single stimulus presentation (gray) of 533 ms. In total, twelve images were extracted, whereby one trial was excluded due to artifacts, resulting in a total of eleven images. Broadband timecourses are averaged across electrodes and visual areas. **B**: The ratio between the transient and sustained response magnitude against two summary variables describing low-level image statistics, including the contrast energy (CE, *left*) and spatial coherence (SC, *right*). Data points depict individual images and the red line depicts a linear regression fit.

